# Crosstalk between glucose metabolism and morphogen signalling specifies tonotopic identity in developing hair cells

**DOI:** 10.1101/2022.04.11.487851

**Authors:** James DB O’Sullivan, Thomas S Blacker, Claire Scott, Weise Chang, Mohi Ahmed, Val Yianni, Zoe F Mann

## Abstract

In vertebrates with elongated auditory organs, mechanosensory hair cells (HCs) are organised such that complex sounds are broken down into their component frequencies along the proximal-to-distal long (tonotopic) axis. Acquisition of frequency-specific morphologies at the appropriate positions along the chick cochlea, the basilar papilla (BP), requires that nascent HCs determine their tonotopic positions during development. The complex signalling within the auditory organ between the developing HC and its local niche along the axis is currently poorly understood. Here we apply NAD(P)H fluorescence lifetime imaging (FLIM) to reveal metabolic gradients along the tonotopic axis of the developing BP. Re-shaping these gradients during development, by inhibiting different branch points of cytosolic glucose catabolism, alters normal morphogen signalling and abolishes tonotopic patterning, normalising the graded differences in hair cell morphology along the BP. These findings highlight a causal link between morphogen signalling and metabolic reprogramming in specifying tonotopic identity in developing auditory HCs.

## Introduction

Hearing relies upon the life-long function of mechanosensory hair cells (HCs) and their associated glial-like supporting cells (SCs) within the cochlea. In both mammals and birds, different frequencies stimulate HCs located at different positions along the basal-to-apical long axis of the auditory epithelium to separate complex sounds into their spectral components. This phenomenon, known as tonotopy, underlies our ability to differentiate between the high pitch of a mosquito and the low rumbling of thunder. The specific factors regulating the development of tonotopy remain, largely, unclear. As high frequency HCs show increased vulnerability to insults, including aging ^1^, noise damage ^2, 3^ and ototoxicity ^4^, awareness of the mechanisms underlying the formation of frequency-specific HC properties is crucial to understanding both acquired auditory defects, HC repair and regeneration. Enhanced knowledge of the pathways that drive specification of HC phenotypes at different frequency positions could identify novel strategies to preserve and restore high frequency hearing loss.

Metabolism, encompassing the complex network of chemical reactions that sustain life (summarised in Figure 1), has emerged as a key regulator of cell fate and differentiation ^5^. Reciprocity between metabolic networks and the epigenome has been extensively studied in models of cancer cell biology and tumorigenesis ^6^. Here, chromatin modifying enzymes (involved in histone acetylation and methylation) that drive cell fate switches rely upon metabolic intermediates as co-factors or substrates, highlighting a link between cell metabolism and transcriptional regulation ^7^. Reprogramming between glycolytic and oxidative pathways has also been reported in developing tissues, including migratory neural crest cells ^8^, the zebrafish otic vesicle ^9^, trophectoderm in the mouse embryo ^10^ and the presomitic mesoderm ^11–14^. Nevertheless, a regulatory role for metabolism has not been explored in the context of cell fate and patterning in developing inner ear epithelia. This is, in part, because the classic biochemical approaches from which our knowledge of metabolism has formed involve the destructive extraction of metabolites from a sample. Probing metabolism in this manner, although valuable, means that any spatial organisation of metabolic pathways in complex tissues is lost. As the cochlea contains multiple cell types, investigating the regulation of their development by metabolism requires experimental approaches capable of interrogating metabolic pathways in live preparations with single cell resolution.

**Figure 1.**
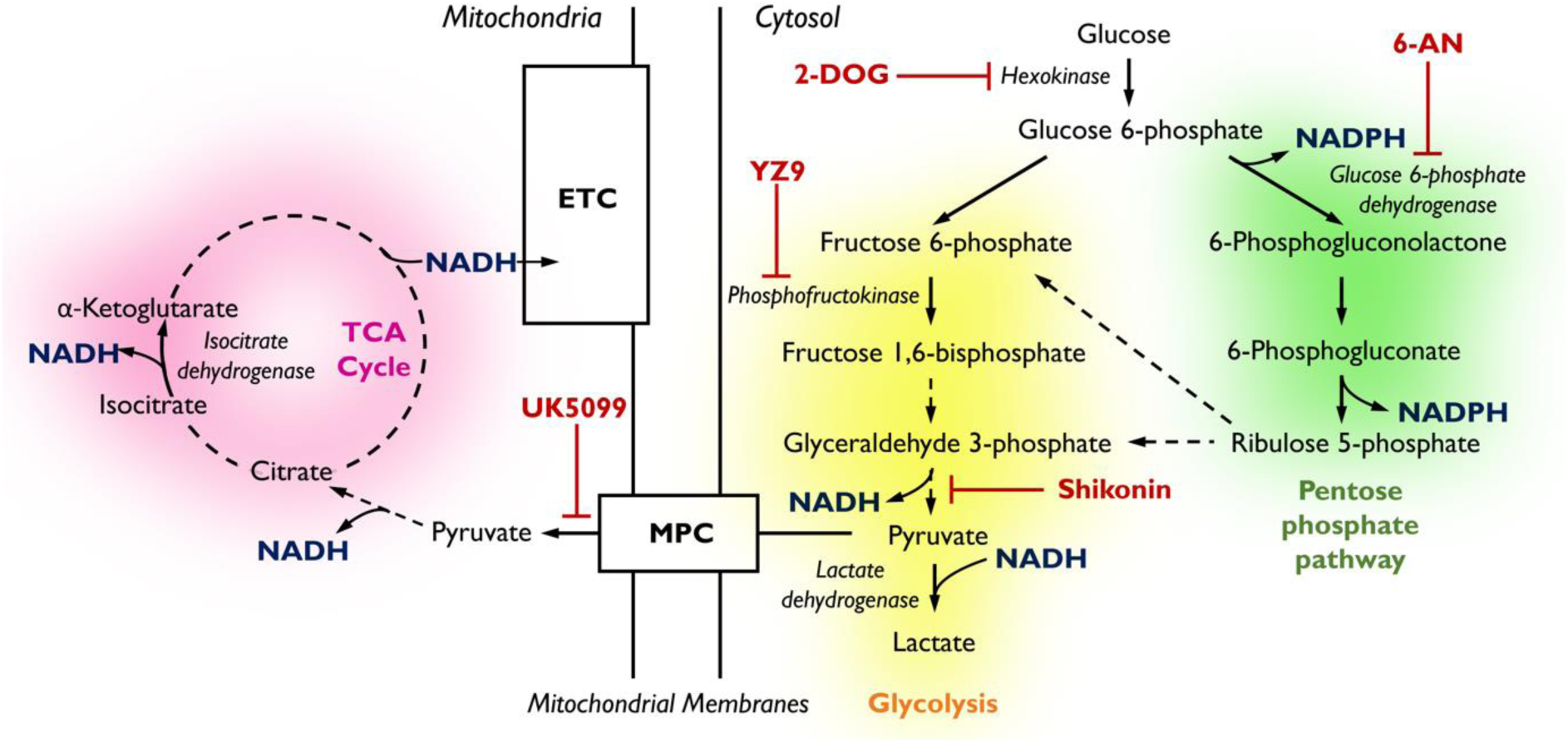
Pathways of glucose catabolism regulating cellular NADPH/NADH. **(pink mitochondrial OXPHOS) - Glucose metabolism in *mitochondria*.** Following its conversion from glucose during glycolysis, pyruvate is transported into the mitochondria via the mitochondrial pyruvate carrier (MPC) and enters the tricarboxylic acid (TCA) cycle. Its sequential oxidation provides reducing equivalents in the form of NADH to the electron transport chain (ETC), driving ATP production by oxidative phosphorylation (OXPHOS). **(yellow glycolysis) - *Cytosolic* glucose flux via the main branch of glycolysis.** In this process, one molecule of glucose is anaerobically converted into two molecules of pyruvate to yield two molecules of ATP. Lactate dehydrogenase (LDH) acts to maintain the pool of NAD+ necessary for glycolysis to take place by oxidising NADH upon the reduction of pyruvate to lactate. **(green pentose phosphate pathway) – *Cytosolic* glucose flux into the oxidative branch of the pentose phosphate pathway (PPP).** Running parallel to glycolysis, the PPP branches off at glucose 6-phosphate (G6P) generating NADPH and ribose 5-phosphate (R5P). PPP shuttles carbons back into the main glycolytic pathway at glyceraldehyde 3-phosphate and fructose 1,6-bisphosphate. Different pathways of glucose flux can be targeted for pharmacological intervention. Inhibitors for various metabolic branch points are indicated in red (UK5099, YZ9, 2-DOG, 6-AN, Shikonin).

We have previously demonstrated that fluorescence lifetime imaging microscopy (FLIM) provides a label-free method to identify metabolic differences between inner ear cell types^15^. By spatially resolving differences in the fluorescence decay of the reduced redox cofactors nicotinamide adenine dinucleotide (NADH) and its phosphorylated analogue NADPH (Figure 1) we can extract information about the metabolic state of a cell ^15^. Here, we apply this technique to investigate a role for metabolism in specifying morphological properties of proximal (high frequency) verses distal (low frequency) HCs in the chick cochlea. The HC phenotypes associated with different tonotopic positions are defined using previously characterised morphometrics as read-outs ^16, 17^. By applying NAD(P)H FLIM in different regions of the developing BP, we identify a gradient in NADPH-linked glucose metabolism along the tonotopic axis. The NAD(P)H gradient did not originate from a tonotopic switch between glycolytic and oxidative pathways or from differences in cellular glucose uptake. We find that the metabolic gradient along the developing BP originates instead from tonotopic differences in the catabolic fate of cytosolic glucose once it has entered the cell. By modulating the flux of glucose through specific cytosolic branches, we systematically interrogated its role in specifying tonotopic properties of developing HCs. We found that the cellular balance of glucose entering the pentose phosphate pathway (PPP) and the main branch of glycolysis (Figure 1) instructs tonotopic HC morphology by regulating the graded expression of Bone morphogenetic protein 7 (Bmp7) and its antagonist Chordin-like-1 (Chdl1), known determinants of tonotopic identity ^18^. This work highlights a novel role for cytosolic glucose metabolism in specifying HC positional identity at the morphological level providing the first evidence of a link between metabolism and morphogen signalling in the developing inner ear.

## Results

### NAD(P)H FLIM reveals differences in the cellular balance between NADPH and NADH along the tonotopic axis of the developing BP

NAD and NADP are metabolic cofactors responsible for ferrying reducing equivalents between intracellular redox reactions throughout the cellular metabolic network (Figure 1). The two molecules are fluorescent in their reduced (electron-carrying) forms NADH and NADPH, a feature that is lost upon oxidation to NAD+ and NADP+. The spectral properties of NADH and NADPH are identical, meaning that their combined signal emitted from living cells is labelled as NAD(P)H. FLIM of NAD(P)H has shown significant promise for identifying changes in the metabolic pathways active at a given location in living cells.

NAD(P)H FLIM typically resolves two fluorescence lifetime components in live cells and tissues one with a duration of around 0.4 ns coming from freely diffusing NAD(P)H (τ_free_) and the other of 2 ns or more from the pool of NAD(P)H that is bound to enzymes and cofactors (τ_bound_) ^19, 20^ (Figure 2B). Changes in the duration of τ_bound_ indicate switching in the enzyme families to which the overall NAD(P)H population is bound. NAD(P)H FLIM can therefore report changes in metabolic state in live cells during physiological processes. We used NAD(P)H FLIM to monitor metabolism along the proximal-to-distal (tonotopic) axis of the BP during development (Figure 2E-H). Around E6, when cells in the BP begin acquiring their positional identity ^18, 21^, we observed a significant difference in τ_bound_ along the tonotopic axis (Figure 2D-H). The proximal-to-distal gradient in τ_bound_ (Figure 2D) was also evident at E9, when a majority of cells are post mitotic ^22^, and at E14, when HCs functionally resemble those in a mature BP (Figure 2E-F). The gradient observed in τ_bound_ throughout BP development, specifically at E6 and E9, is consistent with our previous work, where we showed that graded morphogen signalling along the BP establishes HC positional identity between E6 and E8 ^18, 21^. Changes in τ_bound_ are report shifts in the cellular balance between NADPH and NADH ^15, 23, 24^, which reflects differences in metabolic pathway utilisation (Figure 2 Supplement 1). These data therefore suggest alterations in the balance between NADH- and NADPH-linked metabolism along the tonotopic axis of the developing BP.

**Figure 2.**
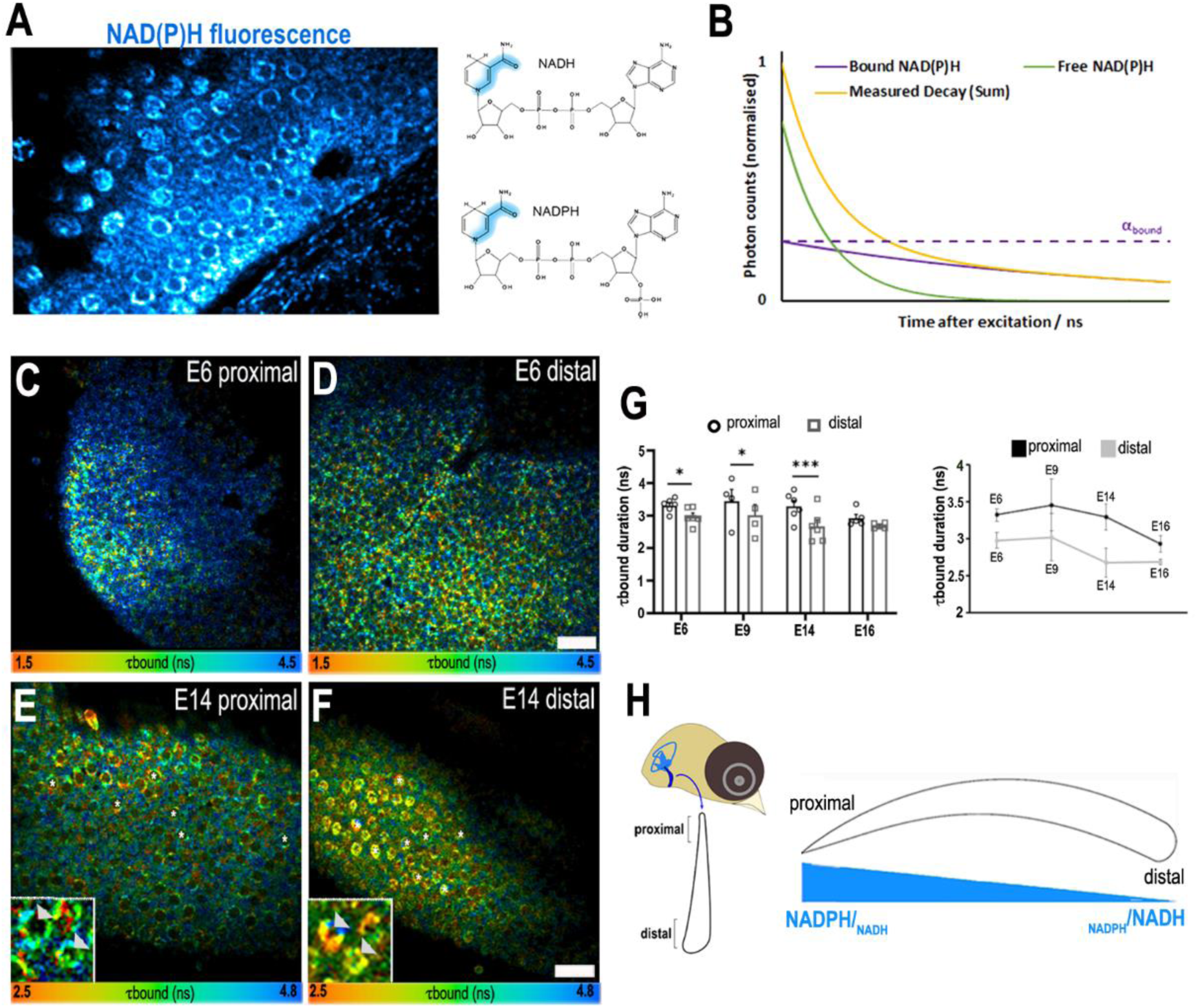
A proximal-to-distal metabolic gradient in the developing chick cochlea. **(A)** Two-photon fluorescence image showing NAD(P)H in a live BP explant at E14 and the origin of inherent fluorescence at the nicotinamide ring. **(B)** NAD(P)H FLIM resolves two components corresponding to freely diffusing (shorter lifetime, τ_free_) and enzyme bound (longer lifetime, τ_bound_). Changes in τ_bound_ imply changes in the specific enzymes to which NAD(P)H is binding and, therefore, the metabolic state of the cell. The proportion of the total NAD(P)H population that is bound to enzymes, labelled α_bound_, determines the relative contribution of the two species immediately after excitation. **(C-F)** FLIM images of the bound NAD(P)H fluorescence lifetime signal τ_bound_ in the proximal and distal BP regions at E6 and E14. White asterisks indicate the HCs. Higher magnification images highlight the differences in τ_bound_ between proximal and distal HCs at E14 (arrowheads). **(G)** Quantification of τ_bound_ during development shows a shift from NADPH to NADH producing pathways. Line graphs highlight differences in τ_bound_ between proximal (black) and distal (grey) BP regions throughout development. Scale bars = 50μm. Data are mean ± SEM; E6: n = 6, E9: n= 4, E14: n = 6 and E16: n = 5 biological replicates. **\*** p < 0.05, *** p < 0.001 2-way ANOVA. **(H)** Schematic of the chick BP, indicating the proximal and distal regions. Proposed gradient in cellular NADPH/NADH and thus glucose flux along the developing BP.

### Live imaging of mitochondrial metabolism and glucose uptake along the tonotopic axis of the developing BP

The redox states of the cellular NAD(P) pools are highly interrelated with the balance between cytosolic glycolysis and mitochondrial oxidative phosphorylation (OXPHOS) ^25, 26^. We therefore tested whether the gradient in cellular NADPH/NADH reflected a progressive shift from glycolytic to mitochondrial OXPHOS, differences in cellular glucose uptake, or in the catabolic fate of glucose by conducting live imaging in BP explants using a range of metabolic indicators. To assess mitochondrial metabolism, we used the potentiometric fluorescent probe tetramethyl-rhodamine-methyl-ester (TMRM), a cell permeable dye that reports mitochondrial membrane potential (ΔΨ_mt_) in living cells ^27^, and is itself a read-out of glycolytically-derived pyruvate oxidation in the mitochondrial tricarboxylic acid (TCA) cycle and the activity of the mitochondrial electron transport chain (ETC) (Figure 1). 2-NBDG is a fluorescent glucose analogue that when transported into cells via glucose (GLUT) transporters provides an estimate of cellular glucose uptake ^28^. Thus, the 2-NBDG fluorescence measured in a given cell after a defined period of loading reflects the rate of glucose uptake by that cell ^28^.

Explants were dual-loaded with 350 nM TMRM and 1 mM 2-NBDG and both fluorescence signals were analysed from single cells between E7 and E16. TMRM fluorescence revealed no significant difference in ΔΨ_mt_ between proximal and distal regions at any developmental stage (Figure 3 A-B). Analysis of TMRM fluorescence revealed a consistently higher ΔΨ_mt_ in fully differentiated HCs compared to SCs from E14 onwards (Figure 3C). To rule out whether the higher TMRM fluorescence occurred due to differences in dye uptake via the HC transduction channel, explants were dual loaded with the permeant MET channel blocker FM1-43 ^29^ (Figure 3 Supplement 1). These findings indicate no significant difference in mitochondrial activity along the tonotopic axis throughout development, suggesting that the gradient in NADPH/NADH reported by τ_bound_ (Figure 2 C-G) does not arise from variations in the balance between cytosolic and mitochondrial ATP production, as often observed in development ^30^ but from differences in the specific route utilised for processing of glucose in the cytosol. This was supported by measurements of 2-NDBG fluorescence in the same cells (Figure 3 Supplement 2A). These analyses revealed no differences in glucose uptake along the tonotopic axis at any developmental stage or between cell types (Figure 3 Supplement 2B, C), suggesting differences in the fate rather than overall flux of glucose underpin the gradient in τ_bound_ (i.e., NADPH/NADH).

**Figure 3.**
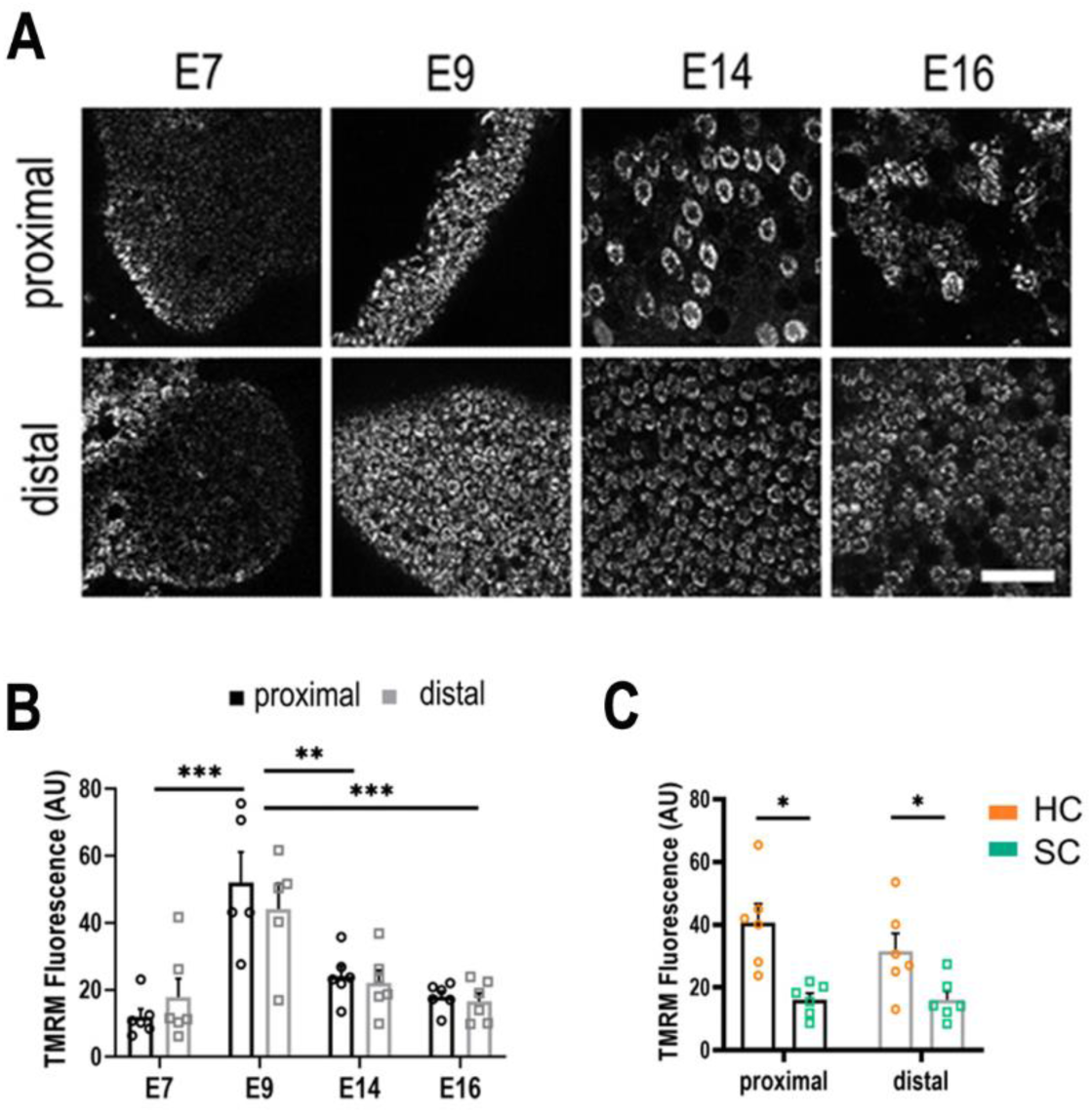
Live imaging of mitochondrial metabolism in HCs and SCs at different positions along the tonotopic axis. **(A)** Mitochondrial membrane potential measured using TMRM in single z-planes from image stacks in the proximal and distal regions of live BP explants. **(B)** TMRM fluorescence indicates a significant increase in mitochondrial activity between E7 and E9, followed by significant decrease between E9 and E14. **(C)** Differences in mitochondrial activity (TMRM fluorescence) between HCs and SCs along the tonotopic axis at E14. Data are mean ± SEM. ** = p > 0.01, *** = p <0.001 for proximal and distal regions 2-way ANOVA. E7 n=6, E9 n=5, E14 n=6, E16 n=6 biological replicates. HCs vs SCs n = 6 biological replicates * = p > 0.05 2-way ANOVA. Scale bar = 40 μm.

### Tonotopic expression of metabolic mRNAs along the proximal-to-distal axis of the developing BP

To further probe the biochemical basis for the gradient in τ_bound_ we exploited existing transcriptional data sets generated from proximal and distal regions of the developing BP ^18^. Prior to mRNA isolation for bulk RNA-seq and Affymetrix microarray analysis, BPs were separated into proximal, middle, and distal thirds. Data were then analysed for differential expression of metabolic mRNAs involved in NADPH regulation and cytosolic glucose flux at E6.5 and E14 (Figure 3 Supplement 3A) ^18^. From the combined data sets, we identified multiple genes involved in NADPH-linked glucose metabolism with differential expression along the tonotopic axis (Figure 3 Supplement 3A). Expression of these genes was verified using RNA scope and Immunohistochemistry. The only enzyme linked with cytosolic glucose flux and cellular NADPH/NADH showing a consistent differential expression throughout development, using all three validation methods, was pyruvate kinase M2 (PKM2) (Figure 3 Supplement 3A, Figure 4 Supplements 1-3)**. No porbe controls were used to validate labelling (Figure 3 Supplement 5).**

**Figure 4.**
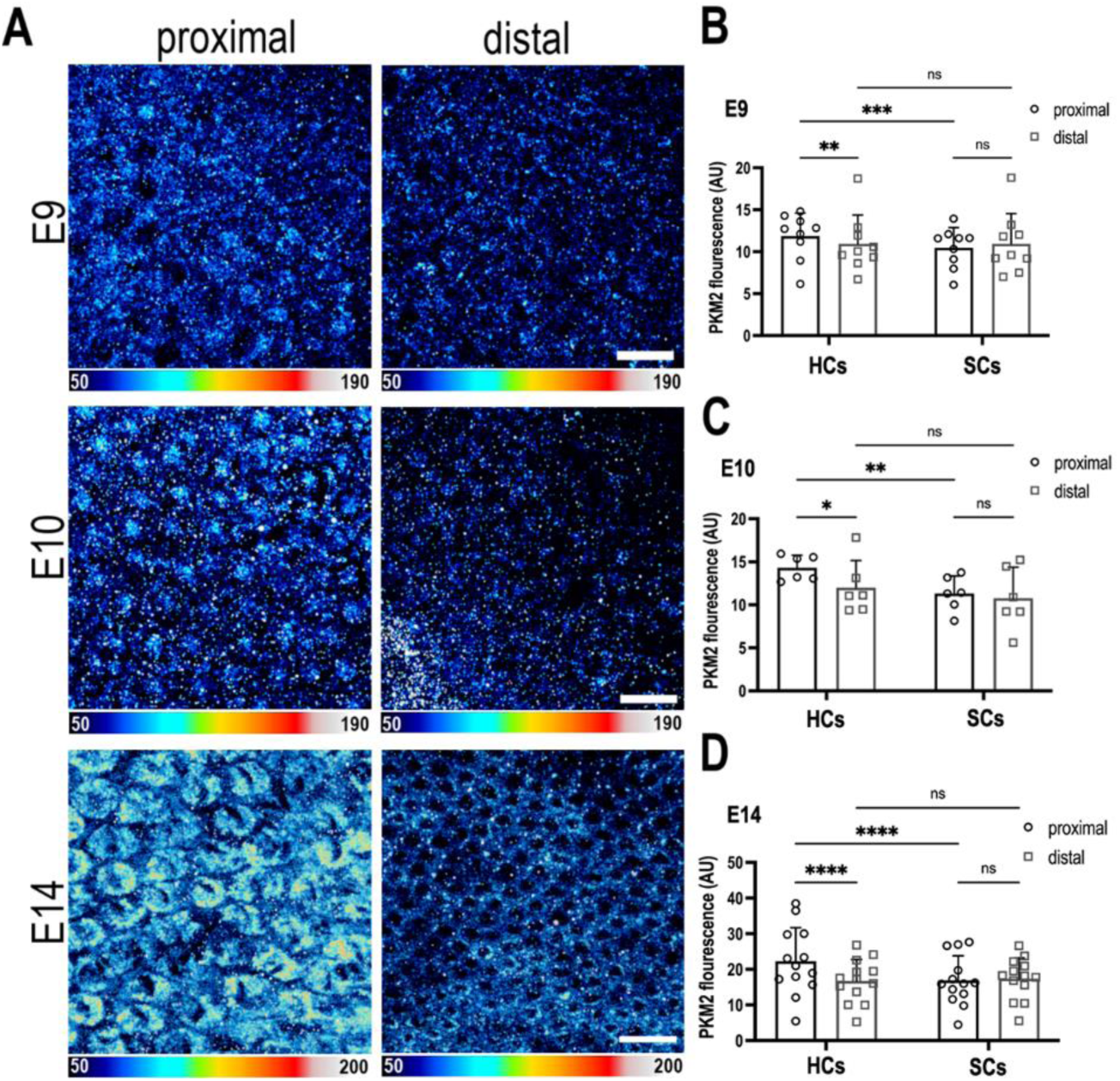
The metabolic gatekeeper Pkm2 is expressed in a Tonotopic gradient during BP development. **(A)** BP whole-mounts labelled for the metabolic enzyme Pkm2 in proximal and distal regions throughout development (E9, E10 and E14). Images show Pkm2 expression at the level of the HC nuclei. Epithelial z-position was determined using Phalloidin and Calbindin staining within the same preparation (images not shown). **(B-D)** Quantification of Pkm2 fluorescence intensity in proximal and distal BP regions at E9, E10 and E14. HC and SC ROIs were determined using Phalloidin and Calbindin staining within the same preparation. Data are mean ± SEM. E9: n = 9, E10: n = 6, E14: n = 13 independent biological replicates. * p = < 0.05, ** p = < 0.01, *** p = < 0.001, p = <0.0001 2-way ANOVA. Scale bars are 20 μm.

### PKM2 protein is expressed tonotopically along the developing BP

Pyruvate kinase M2 (PKM2) is a unique splice isoform of the enzyme pyruvate kinase (PK) and catalyses the final rate-limiting step in glycolysis. PKM2 regulates activity of metabolic enzymes in the upper branch of glycolysis by acting as a gatekeeper and diverting glucose flux towards pyruvate production or into the PPP ^31^. Given this regulatory role in the catabolic fate of cytosolic glucose ^31^ and, the fact that increased PKM2 activity is linked with higher cellular NADPH ^32^, we hypothesised that in correlation with the τ_bound_ gradient, PKM2 expression would be higher in the proximal BP region. Consistent with the observed gradient in NAD(P)H (Figure 2) and in PKM2 mRNA levels (Figure 3, Supplement 3A, Figure 4 Supplements 1-3), we show higher PKM2 protein expression in HCs but not SCs at the proximal compared to distal end of the BP between E9 and E14 (Figure 4).

### Higher intracellular pH in the proximal BP favours PKM2 activity associated with an increased NADPH/NADH ratio and glucose flux into the PPP

The metabolic function of PKM2 is determined by whether the enzyme exists as a tetramer or a dimer. In its dimeric form, PKM2 functions as a metabolic switch, diverting glucose towards the PPP for biosynthesis or towards pyruvate for energy production. ^33^. Allosteric modifications regulating the ratio between the tetrameric and dimeric forms of PKM2 are driven by factors in the surrounding environment including intermediate metabolites and pH ^33, 34^. Given the pH-dependent nature of PKM2 allostery and that the main rate-limiting enzymes driving PPP-linked glucose metabolism display optimal activity at alkaline cytosolic pH ^35^, we next investigated differences in intracellular pH (pH_i_) along the tonotopic axis using the indicator pHrodo Red. When using this probe, low pHrodo Red fluorescence reflects an alkaline pH and high fluorescence a more acidic pH.

Explants were dual-loaded with the pH_i_ indicator pHrodo Red (Figure 5) and the live probe SIR-actin to distinguish HCs from SCs (Figure 5 Supplement 1, 2). We identified opposing proximal-to-distal gradients in pH_i_ in HCs and SCs along the tonotopic axis, using pHrodo Red, which reported a more alkaline pH_i_ in HCs at the proximal compared to distal end of the organ (Figure 5). The higher pH_i_ in the proximal region reflects a metabolic phenotype consistent with higher PPP activity and dimeric PKM2. Overall, the higher pH and PKM2 expression levels and the possible dimeric confirmation are consistent metabolically with a longer τ_bound_ (NAD(P)H lifetime). To investigate whether the proximal-to-distal gradient in pH was maintained at later developmental stages, we also quantified the pHrodo Red signal in HCs and SCs at E14. At later developmental stages, we find the pH gradients to be reversed (Figure 5 Supplement 3). As tonotopic patterning and positional identity are specified between E6-E7.5 ^18^ the gradient at E14 is unlikely to impact the gradient in HC morphology.

**Figure 5.**
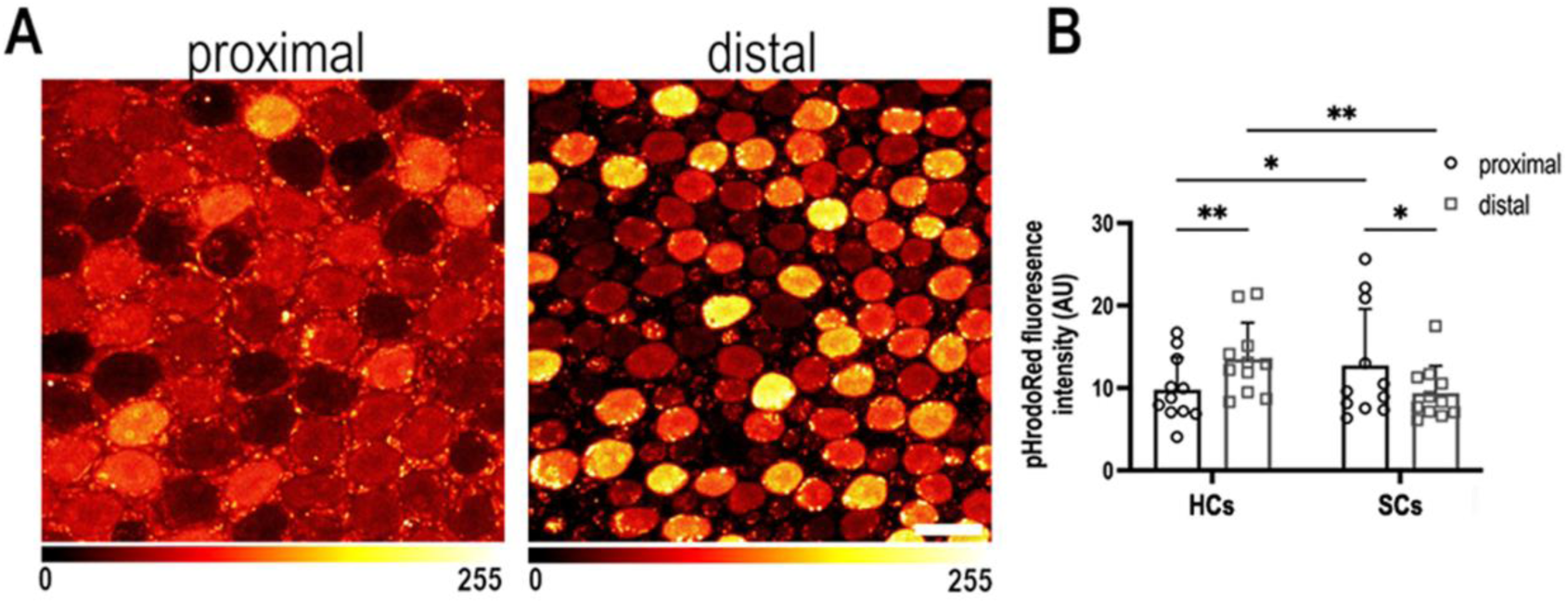
Intracellular pH varies as a function of frequency position during early BP development. **(A)** Intracellular pH, reported by pHrodo Red fluorescence intensity, in the proximal and distal BP regions at E9. High fluorescence indicates acidic pH and low fluorescence a more alkaline pH. **(B)** Mean pHrodo Red fluorescence in measured from HCs and SCs in proximal and distal frequency BP regions. Note the proximal-to-distal gradient in intracellular pH. Data are mean ± SEM from 11 independent biological replicates for HCs and 12 independent biological replicates for SCs. * p = < 0.05, ** p = < 0.01 2-way ANOVA. Scale bar is 10 μm.

### Cytosolic glucose metabolism is necessary for tonotopic patterning in the chick BP

Since we could not detect an obvious transcriptional basis for the proximal-distal gradient in NAD(P)H τ_bound_, we sought to investigate a functional role for metabolism in tonotopic patterning by systematically inhibiting glucose flux into different metabolic pathways (Figure 1). First, we blocked the entirety of cytosolic glucose metabolism using 2-deoxy-d-glucose (2-DOG), an inhibitor of the enzyme hexokinase ^36^, which occurs upstream of the branching of PPP and glycolysis. BP explants were established at E6.5 and maintained for 7 days *in vitro* to the equivalent of E13.5 in control medium or that containing 2mM 2-DOG supplemented with 5mM sodium pyruvate, ensuring adequate substrate supply to the TCA cycle. In a normal BP, proximal HCs have larger luminal surface areas and cell bodies and are more sparsely populated compared to those in the distal region ^16, 17^. These morphological gradients are recapitulated in BP explant cultures during development ^18^. Here, these metrics were determined by measuring differences in the HC lumenal surface area, the size of HC nuclei and the HC density within defined ROIs (100 x100 μm^2^) along the length of the organ. Lumenal surface area was measured using Phalloidin staining at the cuticular plate and nuclear size with DAPI.

In control cultures, HCs developed with the normal tonotopic morphologies (lumenal surface area, nuclear size and gross bundle morphology) (Figure 6A, C, D, Figure 6 Supplement 1). In contrast, when glucose catabolism was blocked between E6.5 and E13.5 equivalent, tonotopic patterning was abolished. This was indicated by a uniformly more distal-like HC phenotype along the BP (Figure 6A, B). In addition to changes in HC morphology, treatment with 2-DOG caused a significant increase in HC density in the proximal but not distal BP region (Figure 6 Supplement 2A, C) again consistent with loss of tonotopic patterning along the organ.

**Figure 6.**
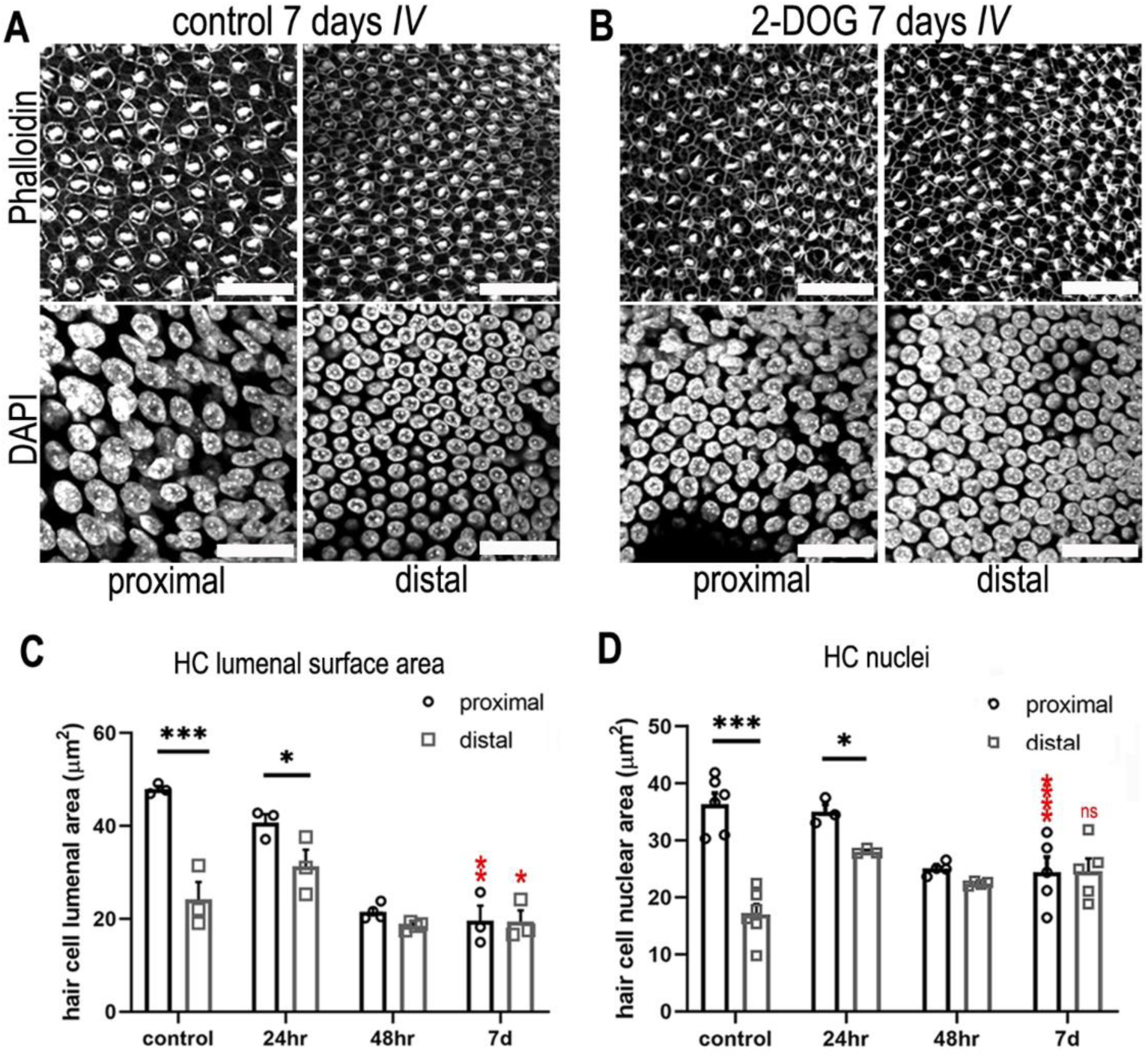
Blocking cytosolic glucose metabolism at key stages of cochlear development induces distal-like phenotypes in the proximal BP. **(A-B)** Maximum z-projections of BP explants showing Phalloidin and DAPI staining in the proximal and distal regions. Explants were maintained from E6.5 for 7 days *in vitro* (equivalent to E13.5) in either control medium or medium supplemented with 2 mM 2-DOG + 5 mM Sodium Pyruvate (NaP). Phalloidin staining depicts differences in HC morphology between proximal and distal regions and DAPI indicates the gradient in hair cell size. **(C)** HC lumenal surface area measured in 2500 μm^2^ ROIs in the proximal (black bars) and distal (grey bars) BP regions for all culture conditions. In controls, mean lumenal surface decreases progressively from the proximal-to-distal region. This gradient is abolished if glucose catabolism is blocked with 2-DOG between E6.5 and E13.5. 2-DOG caused a significant decrease in HC size in the proximal but not distal region. 2-DOG treatments were reduced to 24 or 48 hours to identify the developmental time window during which glycolysis takes effect. Following wash-out of 2-DOG after 24 hours, explants developed with normal HC positional identity. Explants treated with 2-DOG for 48 hours showed no recovery of positional identity following wash-out indicated by the flattening of HC morphology along the BP. **(D)** Quantification of HC nuclei area in the same 2500 μm^2^ ROI areas. Treatment with 2-DOG induced similar, yet less pronounced effects to those seen at the HC cuticular plate. Data are mean ± SEM. **\*** p < 0.05, *** p < 0.001 2-way ANOVA. Controls: n = 6, controls LSA: n = 3; 2-DOG; n = 5, 24 2-DOG, n = 3 and 48h 2-DOG; n = 3 biological replicates. Red stars indicate 2-way ANOVA tests between proximal control and proximal 2-DOG and distal control and distal 2-DOG conditions. To ensure adequate substrate supply to the TCA cycle, 2-DOG-treated explants were supplemented with NaP. G6P glucose 6-phosphate, F6P – fructose 6-phosphate, F16BP – fructose 1,6-bisphosphate, 2-DOG – 2-deoxyglucose. Scale bars are 20 μm.

Changes in glucose metabolism have been linked with reduced cellular proliferation ^37^. We therefore investigated the effects of 2-DOG on proliferation in developing BP explants. We hypothesised that because the majority of cells in the BP are postmitotic by E10 ^22^, adding EdU to cultures in the presence and absence of 2-DOG for 48 hours between E8 and E10 would capture any 2-DOG-dependent differences in proliferative capacity. We observed a consistent reduction in proliferation throughout the whole explant when glucose metabolism was blocked with 2-DOG (Figure 6 Supplement 2B, D).

Increased proliferation is therefore unlikely to account for the higher cell density observed in the proximal region following the inhibition of glycolysis. Further studies are needed to determine the specific mechanisms underlying this frequency-specific increase in HC density.

As shown in our previous work, reciprocal morphogen gradients of Bmp7 and Chdl1 establish HC positional identity at the morphological level along the developing BP between E6.5 and E8 ^18^. To determine whether cytosolic glucose metabolism acts during this same developmental window, we blocked hexokinase activity for defined periods during BP development using 2-DOG. Explants were established at E6.5 and treated for either 24 or 48 hours followed by wash out with control medium. These treatments correspond to the developmental window (E6.5-E8) described previously for refinement of tonotopic morphologies in developing HCs along the proximal-to-distal axis ^18^. The gradient in HC morphology developed normally in BPs treated with 2-DOG for 24 hours but was absent in those treated for 48 hours (Figure 6 C-D, Figure 7A & C, D). These results suggest that glucose metabolism acts within the same developmental time window as Bmp7 and Chdl1 to set up tonotopic patterning along the BP. These findings are also consistent metabolically with the proximal-to-distal gradient in NADPH/NADH (τ_bound_) observed at E6, E9 and E14 (Figure 2G).

**Figure 7.**
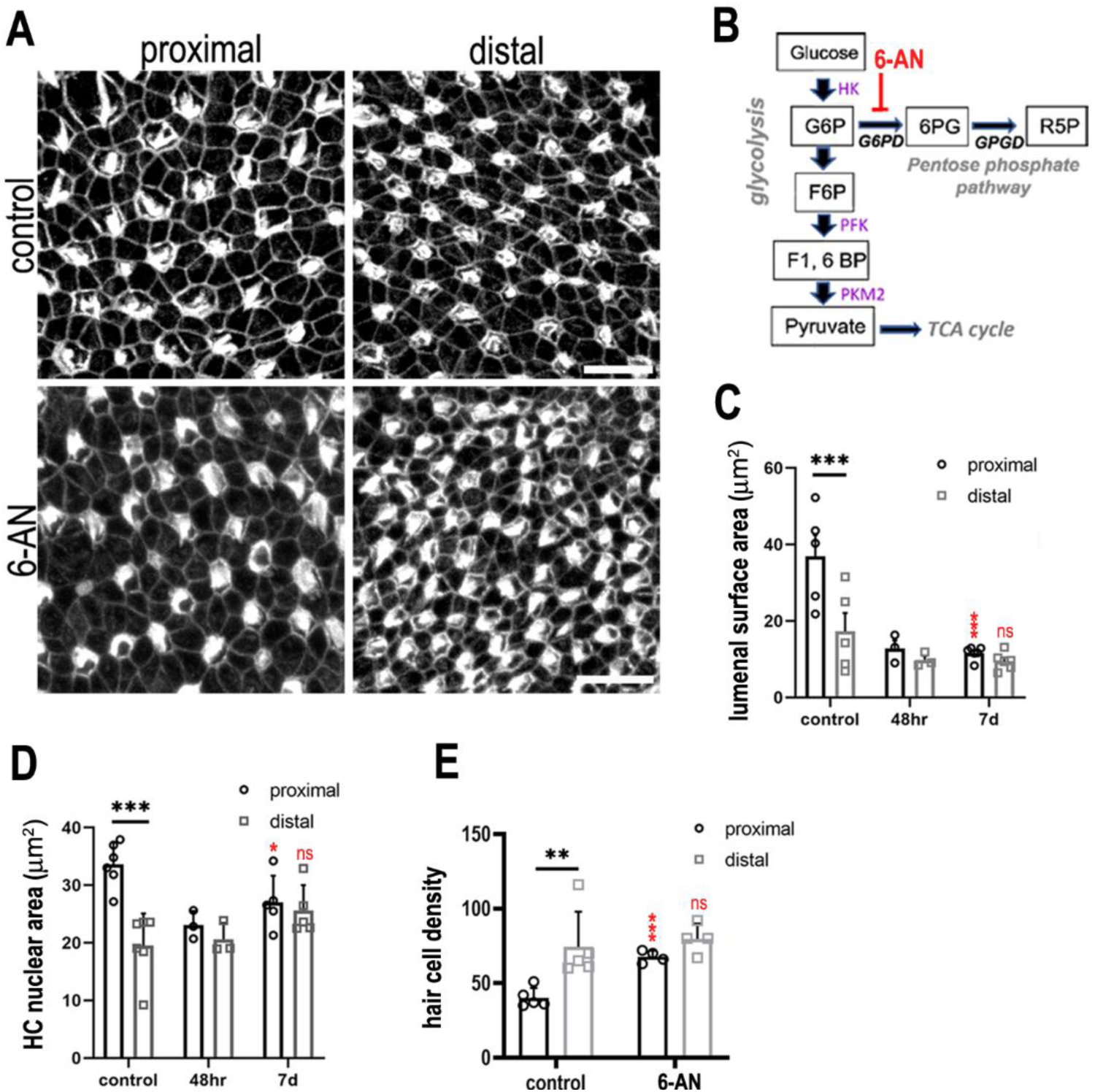
Glucose flux through the pentose phosphate pathway modulates hair cell development and positional identity along the tonotopic axis of the BP. **(A-B)** Maximal z-projections of BP explants cultured from E6.5 for 7 days *in vitro* with control medium or medium containing 2 μM 6-AN. Images show the epithelial surface in proximal and distal BP regions stained with Phalloidin. **(C)**Treatment of explants with 6-AN, a specific blocker of flux through the pentose phosphate pathway, caused a significant reduction in proximal compared to distal hair cell lumenal surface area. **(D-E)** Quantification of hair cell luminal surface area, nuclear area and cell density in defined 2500 μm^2^ ROIs from the proximal (black bars) and distal (grey bars) BP regions. Data are mean ± SEM. ** p = < 0.01, *** p = < 0.001 2-way ANOVA. Nuclei: controls: n = 6, 48h n = 3, 7 days n = 5; LAS: controls n= 5, 48h n = 3, 7 days n = 5; cell density: controls n = 5, 7 days n = 4. Independent biological replicates. Red stars indicate statistical significance for proximal and distal regions when compared between control and 6-AN conditions. Scale bars are 10 μm. G6P – glucose 6-phosphate, F6P – fructose 6-phosphate, F16BP – fructose 1,6-bisphosphate.

To further confirm a role for cytosolic glucose metabolism in establishing HC positional identity, we employed a second method of modulating the pathway independently of hexokinase activity ^38^. The rate of cytosolic glycolysis can be lowered indirectly by raising cytosolic levels of the metabolite s-adenosyl methionine (SAM). Consistent with 2-DOG treatment, explants incubated with SAM between E6.5 and E13.5 lacked correct tonotopic patterning indicated by the flattening of HC morphologies along the tonotopic axis (Figure 6 Supplement 3).

### Flux through the PPP is important for tonotopic patterning along the BP

Studies in other systems have linked both the PPP and TCA cycle with cell fate decisions during development and differentiation ^10, 39^. We therefore sought to further dissect the metabolic signalling network during specification of tonotopy in the developing BP. To investigate a role for PPP-linked glucose metabolism, BP explants were established as described, and treated with 50 mM 6-Aminonicotinamide (6-AN) between E6.5 and E13.5 equivalent. Treatment with 6-AN inhibits the rate-limiting PPP enzyme glucose-6-phosphate dehydrogenase (Figure 1). By comparison with control cultures, inhibition of PPP metabolism caused a significant decrease in hair cell size within 2500 μm^2^ areas measured in the proximal BP (Figure 7, Figure 7 Supplement 1).

To determine whether this effect was specific to glucose flux through the PPP we also blocked phosphofructokinase (PFK), a rate limiting enzyme further down in the glycolytic pathway, using 1 mM YZ9 (Figure 7 Supplement 2). Blocking PFK activity inhibits the glycolytic cascade involved in pyruvate production but does not change the activity of G6PD in the PPP ^10^. YZ9 treatment resulted in a reduction in HC size, especially in the proximal region leading to a reduction in the HC gradient when analysed using pairwise comparisons (Sidak’s multiple comparisons, p=0.17). However, contrary to all other metabolic inhibitor treatments, YZ9 was unique in that it did not produce a significant interaction term in our 2-way ANOVA (Figure 7 Supplement 2C, Supplementary Table 1). The interaction term indicates detection of differences in the proximal-distal gradient that are induced by YZ9 treatment relative to the control group. These analyses demonstrates that YZ9 is significantly less disruptive to the proximal-distal gradient than 2-DOG and SAM, which inhibit both glycolysis and the PPP, and 6-AN, which is PPP specific (Figure 7 Supplements 1,2, Supplementary Table 1). These findings suggest that tonotopic patterning and specification of HC positional identity are regulated by glucose flux into the PPP, rather than by enzymes such as PFK in the lower branch of glycolysis, consistent with the higher NADPH/NADH ratio in these regions observed using FLIM.

### Combined activity in PPP and PKM2 pathways are required for specification of HC positional identity and patterning in the developing BP sensory epithelium

Having identified graded differences in Pkm2 expression (Figure 3 Supplement 3, Figure 4), we investigated whether the enzyme is required for specification of HC morphologies during development. Pkm2 activity was blocked during development from E6.5 to E13.5 equivalent using the pharmacological inhibitor shikonin (Figure 8). Treatment of explants with shikonin during HC formation abolished the normal tonotopic gradient in HC lumenal surface area (Figure 8A, C), where HC cuticular plate circumference was significantly increased in the distal but not proximal BP region compared to controls. Absence of Pkm2 activity did not alter HC nuclear area (Figure 8D), suggesting the enzyme could play a more important role regulating HC morphology and patterning at the epithelial surface.

**Figure 8.**
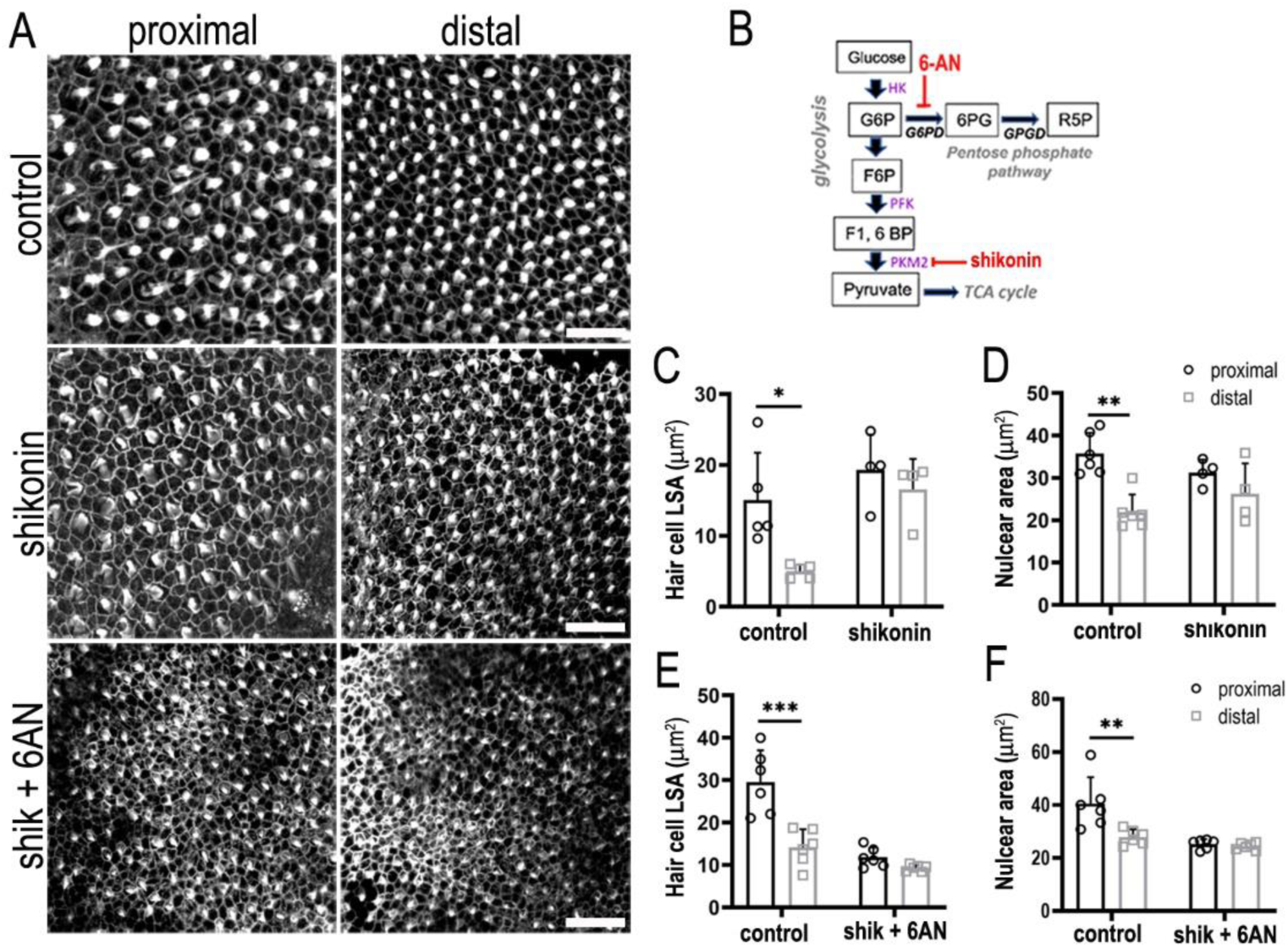
Glucose catabolism in PPP and Pkm2 pathways is necessary for specifying correct HC morphology and patterning along the tonotopic axis of the developing BP. **(A)** Maximal z-projections showing Phalloidin staining at the epithelial surface in proximal and distal BP regions of control and shikonin and shikonin + 6-AN-treated explants. **(B)** Schematic illustrating the metabolic enzymes targeted by shikonin and 6-AN. **(C-D)** Quantification of HC lumenal surface area and nuclear are in control and shikonin BP explants established at E6.5 and maintained for 7 days *in vitro*. **(E-F)** Quantification of HC lumenal surface area and nuclear are in control and shikonin + 6-AN explants treated from E6.5 for 7 days *in vitro*. Hair cell circumference, nuclear area and HC density were quantified in 2500 μm^2^ ROIs in proximal and distal regions. Data are mean ± SEM. * p = < 0.05, ** p = < 0.01, *** p = < 0.001 2-way ANOVA. Controls: n = 5 for LSA, n = 6 for nuclear area, shikonin: n = 4 shikonin + 6-AN: n = 6. Scale bars are 20 μm.

2-DOG blocks the activity of all pathways in cytosolic glycolysis, a network that encompasses both Pkm2 and PPP-linked glucose catabolism. Having probed Pkm2 and PPP metabolism independently we tested whether blocking both pathways during development was additive and could mimic the inhibitory effects of 2-DOG on HC positional identity (Figure 6). We find that blocking both pathways during development abolished the normal gradient in HC morphology along the tonotopic axis thus mimicking the effects seen when inhibiting glycolysis using 2-DOG (Figure 8A, E, F). These data therefore suggest that metabolic activity in both PPP and Pkm2 pathways is important for establishing HC positional identity along the developing BP.

### Pyruvate-mediated OXPHOS in mitochondria maintains progression of HC development but does not regulate tonotopic patterning

In the developing BP, live imaging of mitochondrial activity using TMRM revealed no apparent difference in OXPHOS along the tonotopic axis. To determine whether mitochondrial metabolism influences tonotopic patterning during development, we blocked uptake of glycolytically-derived pyruvate into mitochondria by inhibiting the mitochondrial pyruvate carrier (MPC) with UK5099 ^40^ (Figure 9B). HCs in explants treated with UK5099 between E6.5 and E13.5 developed with abnormal morphologies at all positions along the BP and displayed immature stereociliary bundles or lacked them completely (Figure 9A, C, red arrows). To determine whether this effect was due to a global developmental arrest, explants were also established at E8, by which time tonotopy is specified ^18^ but bundles are not yet developed, and maintained for 7 days *in vitro* to the equivalent of E15. Compared to controls, HCs of explants treated with UK5099 were smaller and displayed abnormal bundle morphologies at all positions along the BP (Figure 9C, Figure 9 Supplement 1). The role that mitochondria play in shaping HC morphologies and functional properties at different frequency positions is at present unclear and will require further investigation. However, our findings suggest that pyruvate mediated OXPHOS plays a more significant role in maintaining the overall progression of development rather than regulating patterning along the tonotopic axis.

**Figure 9.**
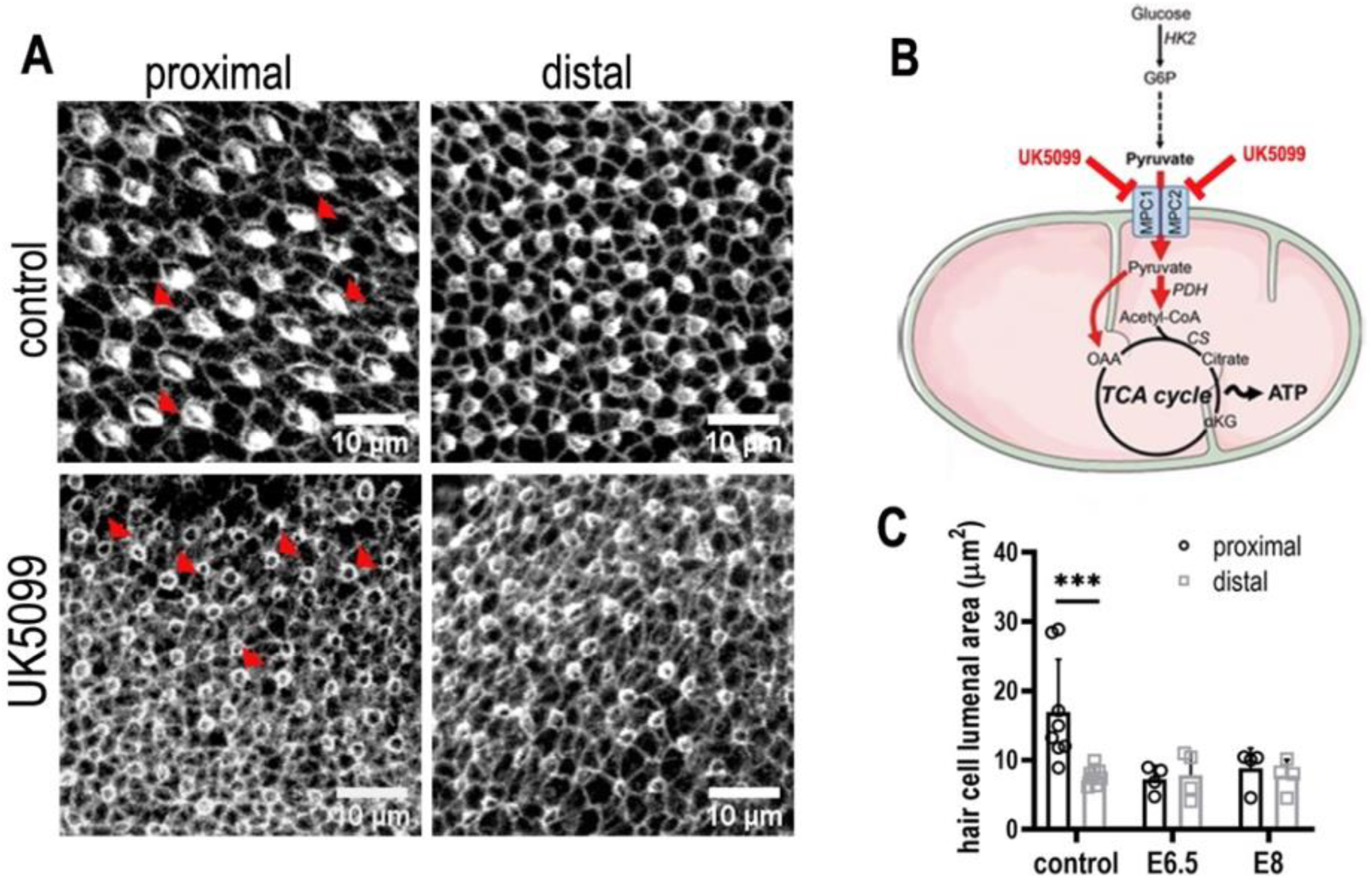
Mitochondrial OXPHOS is necessary for the normal developmental progression of hair cells but not positional identity. **(A)** Hair cell morphology at the surface of the BP epithelium in explants stained with Phalloidin. Cultures were established at E6.5 and maintained for 7 days *in vitro* in control medium or that supplemented with the mitochondrial inhibitor UK5099. **(B)** Blocking pyruvate uptake into mitochondria with UK5099 disrupts normal TCA cycle activity and thus mitochondrial OXPHOS by blocking pyruvate uptake via the mitochondrial pyruvate carrier (MPC1/MPC2). **(C)** Blocking mitochondrial OXPHOS from E6.5 to E13.5 equivalent caused developmental abnormalities in HCs along the BP including reduced HC size and immature stereocilial bundles (red arrowheads in A) in both proximal and distal regions compared to controls. To test whether mitochondrial OXPHOS is required for overall developmental progression, cultures were also established at E8 and treated with 1μM UK5099 for 7 days *in vitro* to the developmental equivalent of E15. HCs showed no apparent recovery of normal morphology following UK5099 treatment from E8 compared to E6.5. Data are mean ± SEM. *** p < 0.001, 2-way ANOVA. Controls: n = 8, UK5099 E6.5: n = 4, E8: n = 4.

### Glucose metabolism regulates expression of *Bmp7* and *Chdl1* along the tonotopic axis

In many developing systems, gradients of one or more morphogen act to regulate cell fate, growth and patterning along a given axis ^41, 42^. In the chick cochlea, reciprocal gradients of *Bmp7* and its antagonist *Chdl1* play key roles in determining HC positional identity. As disruption of both the normal gradient in *Bmp7* ^18^ and glucose metabolism induce similar defects in morphological patterning (Figure 6, Figure 10A-D), we investigated the possibility of a causal interaction between the metabolic and the morphogen signalling networks in the developing BP.

**Figure 10.**
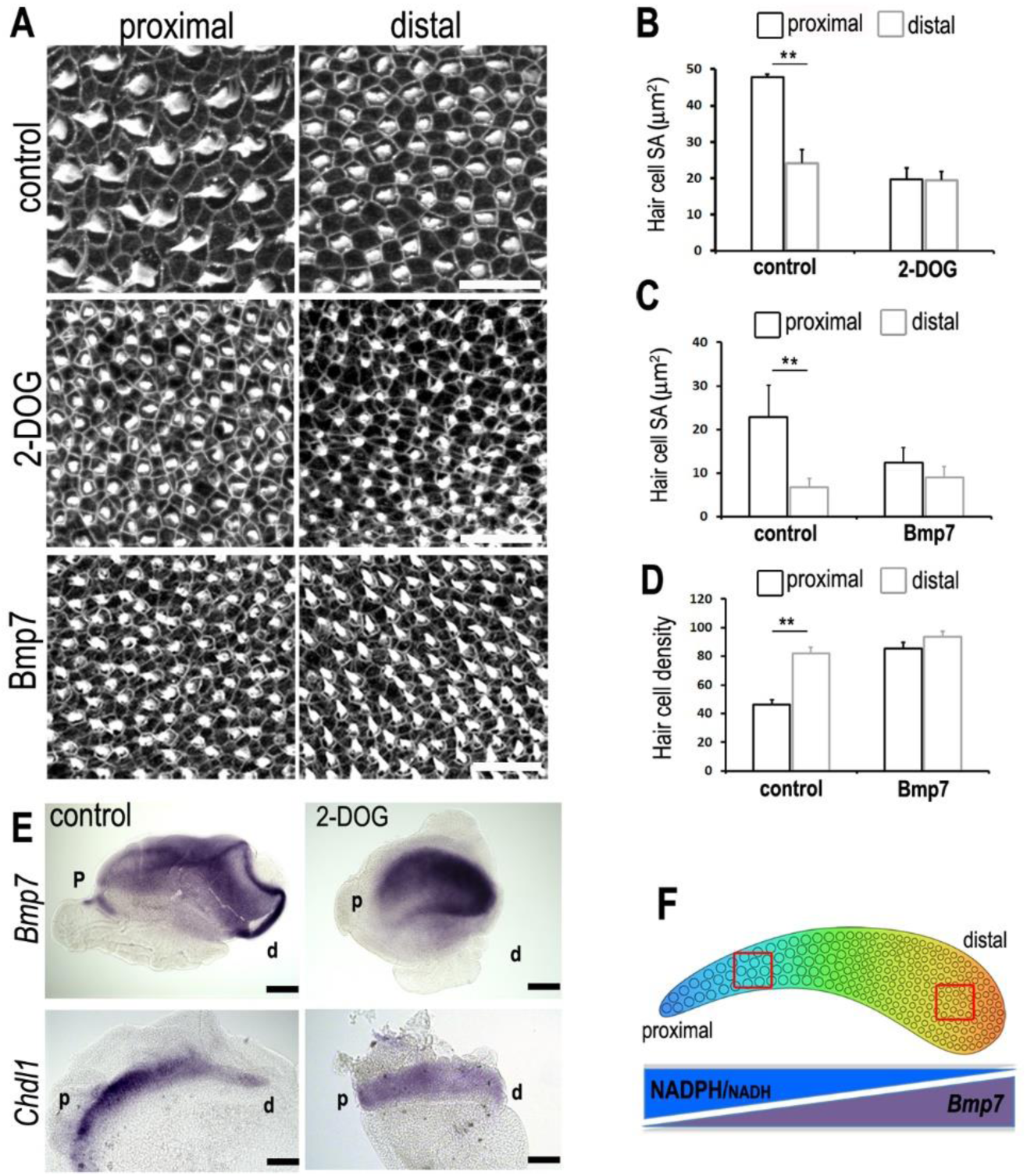
A tonotopic gradient in NAD(P)H producing glucose metabolism specifies hair cell positional identity along the BP by regulating gradients of *Bmp7* and *Chdl1*. (**A**) Phalloidin staining at the surface of BP explants in the proximal and distal regions. Explants were established at E6.5 and incubated for 7 days *in vitro* in control medium or medium containing 2-DOG + NaP or Bmp7. **(B-C)** Treatment with 2-DOG or Bmp7 induced HC morphologies consistent with a more distal phenotype in the proximal BP. Hair cell lumenal surface area was determined using Phalloidin staining at the cuticular plate in 2500 μm^2^ areas. **(D)** Treatment with Bmp7 between E6.5 and E13.5 equivalent results in increased hair cell density in the proximal BP region. Hair cell density was counted in proximal and distal BP regions using defines ROIs of 10000 μm^2^. **(E)** Treatment of explant cultures with 2-DOG + NaP from E6.5 for 72 hours *in vitro* disrupts the normal tonotopic expression of *Bmp7* and its antagonist *Chdl1*. Images show *in situ* hybridisation for *Bmp7* and *Chdl1* in BP whole-mounts treated with 2-DOG + NaP from E6.5 for 72 hours *in vitro*. Images are representative of 6 biological replicates. (2-DOG) Controls: n=6, 2-DOG: n=6 Data mean ± SEM. ** p < 0.01 2-way ANOVA. (Bmp7) Controls: n=11 Bmp7 n=10 Data mean ± SEM. ** p < 0.01 2-way ANOVA. Scale bars (A) control scale bar is 20 μm, DOG and Bmp7 are 50 μm. Scale bars for *in situ* data (E) are 10μm, **(F)** Schematic of the chick BP, showing the graded differences in hair cell size and density along the tonotopic axis. The opposing gradients in *Bmp7* activity and in cellular NAD(P)H/NADH (glycolysis) are indicated. Red boxes indicate regions of measurement for HC lumenal surface areas and cell density.

To investigate the regulatory effects of cytosolic glucose metabolism on the expression gradients of *Bmp7* and *Chdl1*, explants were established at E6.5 and maintained for 72 hours *in vitro* (equivalent of E9.5) in control medium or that containing 2-DOG + sodium pyruvate. Whole-mount *in situ* on explant cultures showed that disrupted glucose metabolism altered the normal expression of *Bmp7* and *Chd1*along the BP (Figure 10E). Following treatment with 2-DOG, *Bmp7* expression appeared to increase along the entire BP while *Chdl1* showed a reciprocal decrease along the length of the organ.

It is challenging to predict how this change in global change in expression levels would impact the activity of each morphogen along the tonotopic axis but it does support our hypothesis that there is a causal interaction between glycolytic and Bmp7-Chdl1 networks. The precise nature of this interaction requires further investigation. We speculate that the increased *Bmp7* and reduced *Chdl1* expression in the proximal region (Figure 10E, Figure 10 Supplement 1), in response to perturbed glucose flux (by treating with 6-AN), would induce expansion of distal-like HC morphologies into the proximal region. Blocking mitochondrial-linked glucose catabolism with UK5099 did not alter the expression of *Bmp7* (Figure 10 Supplement 2).

### Treatment with Chdl1 restores normal tonotopic patterning when glycolysis is blocked during development

Modulating the reciprocal gradients of *Bmp7* and *Chdl1* along the proximal-to-distal axis alters tonotopic patterning in nascent HCs ^18^. We further show that the normal gradients of *Bmp7* and *Chdl1* are disrupted along the BP when glycolysis or PPP activity are blocked during development (Figure 10E, Figure 10 Supplement 1). Treatment with 2-DOG causes a global increase in *Bmp7* and decrease in *Chdl1* expression (Figure 10E), we therefore hypothesised that treatment of explants with 2-DOG in the presence of Chdl1 protein might restore tonotopic patterning when glycolysis is blocked. Analysis of explants treated between E6.5 and E13.5 equivalent with 2-DOG + 0.4 μg/mL Chdl1 showed a partial rescue of HC morphologies (lumenal surface area and nuclear area) along the proximal-to-distal axis (Figure 11, Figure 11 Supplement 1 and Supplementary Tables 1, 2). Whilst the precise length and number of stereocilia could not be accurately quantified using Phalloidin staining in these explants, the overall bundle morphology also appeared consistent with that reported previously for proximal and distal BP regions ^21, 43, 44^.

**Figure 11.**
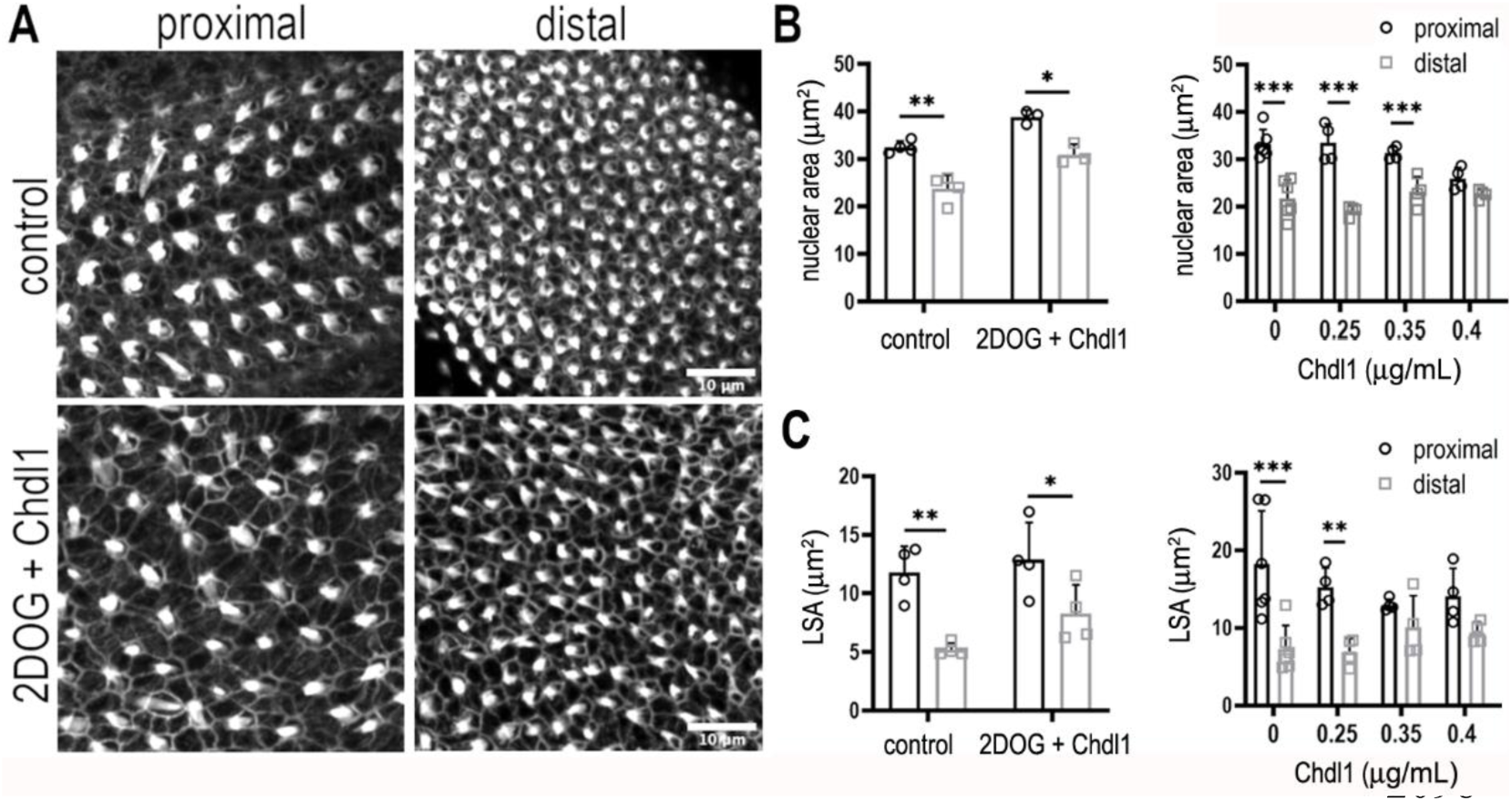
Chdl1 restores tonotopic patterning along the BP when cytosolic glycolysis is blocked with 2-DOG. **(A)** Maximum z-projections of the epithelial surface in the proximal and distal regions of Phalloidin-stained BPs. Explants were established at E6.5 and maintained for 7 days *in vitro* in either control medium or that containing 2-DOG and Chdl1. Phalloidin staining indicates differences in HC lumenal surface area and gross bundle morphology between proximal and distal regions. **(B)** The tonotopic gradient in HC lumenal surface was restored when explants were treated with 2-DOG in the presence of Chdl1. **(C)** The gradient in HC nuclear area was maintained along the proximal-to-distal axis when explants were treated with 2DOG in the presence of Chdl1. Effects of Chdl1 were only apparent at concentrations of 0.35 μg/mL or above. **(D)** HC lumenal surface area measured from 2500 μm^2^ ROIs in proximal (black) and distal (grey) BP regions. Effects of different Chdl1 concentrations on the HC luminal surface are indicated. Data are mean ± SEM. * p < 0.05 2-way ANOVA. Dose response data - Controls: n = 6, Chdl1 0.25 n=4, 0.35 n=4, 0.4 n=4. Chdl1 +2DOG: Controls n=6, Chdl1+2DOG n = 4 biological replicates for HC lumenal surface areas and n = 3 for nuclear area. Scale bars are 10μm.

Taken together, our findings suggest that a distinct metabolic state coupled with a specific morphogen level can regulate HC morphology at different positions along the tonotopic axis during development. These data also provide further evidence indicating a causal interaction between metabolic and morphogen signalling networks during development. Ascertaining a role for cytosolic glucose metabolism in specifying proximal verses distal HC fate, specifically related to frequency tuning, would require a detailed analysis of HC physiological properties. Future work should therefore determine whether altering metabolism affects the developmental acquisition of not only HC morphology, but also the intrinsic electrophysiological properties and firing characteristics documented for HCs at different tonotopic positions ^45–47^.

## Discussion

To successfully address hearing and balance deficits following damage, in aging or after ototoxic insult, we must be able to generate new, correctly functioning HCs from within the sensory epithelium at appropriate positions along the frequency axis. Generating new HCs that recapitulate the features of those in a healthy cochlea requires a detailed knowledge of the cell biology driving their formation. As high frequency HCs are more vulnerable to insult, there is a need to understand the specific factors and signalling pathways that specify different HC subtypes. Over recent years, metabolism has emerged as a key driver of cell fate and function across various biological systems and contexts ^48, 49^. Taking both descriptive and experimental approaches, we characterised regional differences in metabolism along the developing chick cochlea and explored a role for coordinated signalling between known developmental pathways and glucose metabolism in the establishment of HC positional identity. We identify a tonotopic gradient in cellular NADPH, originating from differences in glucose metabolism between high and low frequency HCs. Tonotopic differences in the catabolic fate of glucose in glycolysis or the PPP modulates Bmp7 and Chdl-1 signalling along the developing BP. This study provides the first evidence supporting a role for crosstalk between metabolism and morphogen gradients in the developing auditory system, building on our current understanding of cell fate specification.

### NAD(P)H FLIM reveals a gradient in metabolism along the tonotopic axis of the developing chick cochlea

Using NAD(P)H FLIM, we uncovered a proximal-to-distal gradient in cellular NADPH resulting from tonotopic differences in the fate of cytosolic glucose. The biochemical basis for this gradient was further investigated by exploring differences in mitochondrial acitivty, pH_i_, and the expression of metabolic enzymes along the tonotopic axis. Collectively, these analyses indicate that differences in the fate of cytosolic glucose, ^31, 32, 35^ rather than between glycolytic and oxidative metabolic states underpin the increased τ_bound_ lifetime and higher NADPH/NADH ratio reported here for high frequency HCs. The higher expression of PKM2 and the more alkaline pH identified in proximal HCs are also consistent with a higher cellular NADPH/NADH ratio ^33, 34^.

### PPP metabolism, HC size and positional identity

Cell geometry and size contribute to overall tissue architecture during development are important for long-term function of the cochlea in vertebrates. Cell size, cell membrane composition and metabolic rate are tightly correlated ^50^. PPP-derived NADPH is utilised extensively in proliferating cells during development where it regulates cell cycle progression, differentiation, and growth ^51, 52^. Increased glucose metabolism has been reported in regenerating ^53, 54^ and developing systems ^10, 11, 13, 14^ where it plays an important role in regulating cell fate, behaviour and shape. However, the chosen path of glucose catabolism is context-dependent and differs across processes and between tissues. PPP metabolism regulates cell division and proliferation through its ability to generate lipid and nucleotide precursors ^55^. Increased glucose flux into the PPP but not the main branch of glycolysis was also recently shown to regulate the balance between proliferation and cell death in the regenerating limb^54^. In the BP, HCs exit the cell cycle in three progressive waves following a centre-to-periphery progression beginning at E5. During this process, both HCs and SCs are added in apposition, to the edges of the band of post-mitotic cells that preceded them ^22^. HC differentiation then begins in the distal portion of the BP at around E6 and extends proximally along the cochlea expanding across the width of the epithelium ^17, 56, 57^. These graded differences in HC size along the organ are an essential requirement for correct auditory coding ^47^. As larger cell size is correlated with increased G6PDH activity and thus more glucose flux into the PPP ^52^, the higher activity in the proximal BP region may underlie the graded differences in HC size that arise during development. It could be argued that the smaller cell size induced following metabolic perturbation in the proximal BP is a result of impaired differentiation. However, given that distal HCs maintain their small size in the mature organ, this morphological change observed in the proximal region following treatment with 2DOG and 6-AN is consistent with a distal-like phenotype. Without a detailed characterisation of distal and proximal cell growth from E6 through to adult stages, the puzzle is challenging to resolve.

PPP metabolism is also closely linked with *de novo* synthesis of lipids and cholesterol, which form an integral part of cell membranes ^58^. Functional interactions between ion channel complexes in the membrane and the local lipid environment have been described previously in mammalian ^59, 60^ and avian HCs ^61^. Frequency tuning in the BP relies on the intrinsic electrical properties of the HCs themselves, where graded differences in the number and kinetics of voltage-gated calcium channels (VGCCs) and calcium-sensitive (BK type) potassium channels underlie the ability of HCs to resonate electrically in response to sound ^47^. By regulating cholesterol in the HC membrane, the graded PPP activity observed here may also regulate aspects of HC electrical tuning.

### A causal link between metabolism and morphogen signalling during development sets up HC positional identity

Morphogen signalling gradients have well defined roles in directing cell identity along developing axes, where cells determine their fate as a function of morphogen concentrations at different positions along them ^41, 62, 63^. We showed previously that reciprocal gradients of Bmp7 and Chdl1 establish HC positional identity along the developing BP ^18^. The gradient of Bmp7 is established by Sonic hedgehog (Shh) signalling emanating from ventral midline structures including the notochord and floor plate ^44^. Here we identify a gradient in glucose metabolism that regulates the morphology of developing HCs along the tonotopic axis through a causal interaction with the Bmp7-Chdl1 network. Disrupting this gradient using 2-DOG, SAM or 6-AN mimics the effects on HC morphology reported previously for altered Shh ^44^ and Bmp7 signalling ^18^, which induced distal-like HC phenotypes in the proximal BP. Furthermore, the effects of impaired glycolysis could be partially rescued in explants treated with 2-DOG in the presence of Chdl1. Our findings thus indicate a complex and causal interplay between Bmp7 and Chdl1 morphogen gradients and glucose metabolism in the specification of HC tonotopic identity. Metabolic gradients are also known to regulate elongation of the body axis and somite patterning ^11, 14^. By establishing a gradient in intracellular pH, glycolysis drives graded Wnt signalling and specifies mesodermal versus neuronal cell fate along the developing body axis ^11–13^. The specific role of the pH_i_ gradient along the developing BP is unclear, however given the importance of the Wnt signalling pathway in both cochlear development and regeneration and repair, it will be important to investigate crosstalk between the two in the context of HC formation and tonotopic identity.

In conclusion, our findings indicate a causal link between PPP activity and graded morphogen signalling in specifying HC morphology along the tonotopic axis during development. However, to accurately confirm whether these morphological changes reflect a switch between proximal and distal HC fate a detailed physiological analysis is required. Future work should determine how altering the fate of glucose affects the morphological and functional development of the stereociliary bundle, the intrinsic electrophysiological properties and the firing characteristics of HCs at different tonotopic positions. Untangling further the interactions between components of the Shh, Bmp7 and metabolic signalling networks will advance our understanding of how hair cells acquire the unique morphologies necessary for auditory coding. From what we understand about frequency selectivity in vertebrates ^64^, recapitulation of tonotopy will require that any gradient, and its associated signalling networks, scale correctly in different inner ear sensory patches and across species with varying head size and cochlear lengths. Understanding how the mechanical constraints associated with growth and patterning in different sense organs modulate these networks will advance our understanding of how to drive formation of specific HC phenotypes in inner ear organoid models.

## Materials and Methods

### Embryo care and procedures

Fertilized White Leghorn chicken (*gallus gallus domesticus*) eggs (Henry Stewart & Co. LTD, UK) were incubated at 37.5°C in an automatic rocking incubator (Brinsea®) until use at specific developmental stages between embryonic day 6 (E6) and E16. Embryos were removed from their eggs, staged according to Hamburger and Hamilton (1951) and subsequently decapitated. All embryos were terminated prior to hatching at E21. All procedures were performed in accordance with United Kingdom legislation outlined in the Animals (Scientific Procedures) Act 1986.

### Preparation of BP explants for live imaging studies

BPs were collected from chick embryos between E7 and E16, and explants were established at E13 to E16 in accord with United Kingdom legislation outlined in the Animals (Scientific Procedures) Act 1986. Explants were placed onto Millicell cell culture inserts (Millipore ®) and maintained overnight at 37°C in medium 199 Earl’s salts (M199) (GIBCO, Invitrogen) containing 2% fetal bovine serum and 5 mM HEPES buffer (Life Technologies). For live imaging experiments, cultures were transferred to glass-bottom 50 mm MatTek dishes and held in place using custom-made tissue harps (Scientifica). Cultures were maintained in L-15 medium at room temperature throughout the experiment.

### Basilar papilla culture

Basilar papillae (BPs) were isolated from embryos incubated for between 6 (E6.0) and 8 (E8.0) days and maintained in chilled Leibovitz’s L-15 media (GIBCO, Invitrogen). Papillae were dissected as described previously ^65^ and cultured nerve-side-down on Millicell cell culture inserts (Millipore ®). Cell culture inserts were placed into 35 mm culture dishes containing 1.5 mL of 199 Earl’s salts (M199) medium (GIBCO, Invitrogen) supplemented with 5 mM HEPES buffer and 2% fetal bovine serum (FBS). Papillae were maintained in M199 medium plus vehicle (control media) for up to 7 days *in vitro* (DIV) until the equivalent of E13.5. For all treatments, a minimum of four samples were analysed. The following factors were applied to experimental BPs in culture at the specified concentrations: 2-Deoxyglucose (2-DOG) 2 mM (SIGMA), Sodium Pyruvate (NaP) 5 mM (SIGMA), 6-Aminonicotinamide (6-AN) 2 μM (SIGMA), *S*-(5′-Adenosyl)-*L*-methionine chloride dihydrochloride (SAM) 50 μM (SIGMA), YZ9 1 μM (SIGMA), Shikonin 1 μM (SIGMA), Bmp7 recombinant protein 0.4 μg/mL (R&D Systems 5666-BP-010/CF), Chordin like-1 recombinant protein 0.4 μg/mL (R&D Systems 1808-NR-050/CF). For 2-DOG wash-out experiments, cultures were treated for 24 or 48 hours followed by wash out with control medium for the remainder of the experiment up to 7 days. For paired controls, medium was also changed at 24 and 48 hours in culture. At the conclusion of each experiment (7 DIV), cultures were fixed in 4% paraformaldehyde (PFA) for 20 minutes at room temperature, washed thoroughly three times with 0.1 M phosphate buffered saline (Invitrogen) and processed for immunohistochemistry.

### Fluorescence lifetime imaging

NAD(P)H FLIM was performed on an upright LSM 510 microscope (Carl Zeiss) with a 1.0 NA 40x water-dipping objective using a 650-nm short-pass dichroic and 460±25 nm emission filter. Two-photon excitation was provided by a Chameleon (Coherent) Ti:sapphire laser tuned to 720 nm, with on-sample powers kept below 10 mW. Photon emission events were registered by an external detector (HPM-100, Becker & Hickl) attached to a commercial time-correlated single photon counting electronics module (SPC-830, Becker & Hickl) contained on a PCI board in a desktop computer. Scanning was performed continuously for 2 min with a pixel dwell time of 1.6µs. Cell type (HC vs SC) and z-position within the epithelium was determined prior to FLIM analysis using the mitochondrially targeted fluorescent dye tetramethylrhodamine methyl ester (TMRM). The dye was added to the recording medium, at a final concentration of 350 nM, 45 min before imaging. TMRM fluorescence was collected using a 610±30 nm emission filter. Excitation was provided at the same wavelength as NAD(P)H to avoid possible chromatic aberration. The 585±15 nm emission spectrum of TMRM ensured its fluorescence did not interfere with acquisition of the NAD(P)H FLIM images.

### FLIM data analysis

Following 5 x 5 binning of photon counts at each pixel, fluorescence decay curves of the form ^15^,

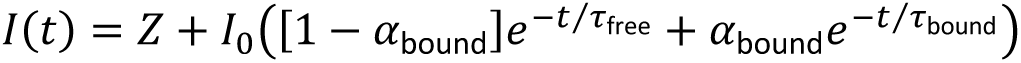

were fit to the FLIM images using iterative reconvolution in SPCImage (Becker & Hickl), where Z allows for time-uncorrelated background noise. Matrices of the fit parameters τ_free_, α_bound_ and τ_bound_ and the total photons counted were at each pixel, were exported and analysed for hair cells and supporting cells, and proximal and distal BP regions, using SPCImage and ImageJ software packages.

### 2-NBDG, TMRM live imaging

BP were isolated from E7, E9, E14 and E16 chick embryos in chilled L-15 mediumand subsequently incubated in 1 mM solution of 2-NBDG (N13195, Thermo Fisher Scientific) in L-15 medium at room temperature for 1 hour. The medium was then replaced with a fresh solution of 1 mM 2-NBDG and 350 nm TMRM (T668, Thermo Fisher Scientific) in L-15 and incubated for a further hour at room temperature. Thereafter, the BPs were washed several times with fresh medium containing 350 nM TMRM and mounted in a 3.5 mm glass bottom MatTek dish. 3D image stacks with an optical thickness of 1 μm were captured using a Leica SP5 confocal microscope with an HCX PL APO 63×/1.3 GLYC CORR CS (21 °C) objective.

### Measurement of pH_i_ using pHrodo Red

BP were dissected in cold L-15 media and incubated for 1 hour at room temperature with 5 µM pHrodo Red Intracellular pH Indicator (Invitrogen P35372) and 1 nM SiR-actin (SpiroChrome SC001) in L-15 medium. Samples were subsequently mounted in Mattek (50 mm) dishes and held in place using custom-made imaging grids. Explants were imaged using an inverted ZEISS LSM980 confocal microscope using a 63X objective and digital zoom of 1.8X. Z-intervals were kept consistent at 0.4 µm across all developmental stages.

### Immunohistochemistry

Inner ear tissue was collected at various developmental stages, fixed for 20 min to one hour in in 0.1 M phosphate buffered saline (PBS) containing 4% paraformaldehyde (PFA), and processed for whole-mounts immunohistochemistry. The BP was then fine dissected and permeabilized in PBS containing 0.3% Triton for 30 min before immunostaining using standard procedures ^18^. Samples were stained with primary antibodies for LDHA 1:75 (Genetex GTX101416) and IDH3A 1:75 (ab228596, Abcam) and Calbindin 1:50 (ab82812, Abcam), PKM2 1:100 (Cell Signalling 4053T). Antibody staining was visualised using secondary antibodies conjugated to either Alexa 488 or Alexa 546 (Invitrogen). Cultures were incubated with all secondary antibodies for 1 hour at room temperature 1:1000, washed thoroughly in 0.1 M PBS. Samples were then incubated for an additional hour with either Alexa Fluor-conjugated Phalloidin-546 1:250 (Invitrogen) to label filamentous actin and DAPI 1:1000 to label cell nuclei. Tissue samples were mounted using Prolong Gold antifade reagent (Invitrogen). 3D image stacks of mounted samples were captured using a Leica SP5 confocal microscope with an HCX PL APO 63×/1.3 GLYC CORR CS (21 °C) objective.

### EdU staining

Control or 2-DOG-treated cultures were incubated for 48 hours in 10 μM 5-ethynyl-2’-deoxyuridine (EdU) from E8 to E10. Cultures were subsequently fixed for 15 minutes in 4% PFA at room temperature and then washed in 0.1 M PBS. Explants were then processed for EdU staining following the Click-iT® EdU 488 protocol (Thermo Fischer Scientific).

### Image analysis

Analysis of z-stacks from immunohistochemistry stains as well as 2-NBDG, TMRM and pHrodo Red live imaging experiments was carried out using the Fiji distribution of ImageJ. For each sample, a z-plane 2 μm beneath the surface of the epithelium was selected using Phalloidin or SiR-actin labelling for further analysis. For each of these selected z-planes, a 100 μm x 100 μm region of interest (ROI) was chosen containing intact tissue in which all HCs were optimally orientated for analysis. Mean fluorescence intensity of the tissue was measured for HCs and SCs from within defined 100 μm x 100 μm ROIs at E7, E9 and E10 timepoints. At E14 and E16, HCs and SCs were manually segmented. At younger stages, when HCs and SCs were not easily identified, fluorescence intensity was measured from within the whole epithelium. HC labels were dilated by 3 μm, which provided selections which included both hair cells and their surrounding SCs. By subtracting the hair cell segmentation from the dilated label, we were thus able to measure the fluorescence intensity of whole HCs separately from their surrounding support cells in the 2-NBDG and LDHA data. A similar approach was adopted when measuring TMRM and IDH3A fluorescence intensity at E14 and E16. However, we noticed that signal was concentrated around the HC periphery. In order to ensure that the fluorescence intensity best reflected only the mitochondria and was not reduced by the low fluorescence from the centre of each HC, we measured mean fluorescence intensity only up to 2 μm from the cell membrane. Likewise, for TMRM and IDH3A data at E7 and E9, mitochondria were segmented using Fiji’s auto-local thresholding (Niblack) prior to intensity measurements, to avoid a biased estimate of fluorescence intensity due to empty space surrounding each mitochondrion.

### Analysis of hair cell morphology

Data were analysed offline using ImageJ software. Hair cell lumenal surface area and cell size were used as indices for HC morphology along the tonotopic axis. To determine the hair cell density, the lumenal surfaces of hair cells and cell size, cultures were labelled with Phalloidin and DAPI. Then, the number of hair cells in 50 μm × 50 μm ROI (2,500 μm^2^ total area) located in the proximal and distal BP regions were determined. Proximal and distal regions were determined based on a calculation of the entire length of the BP or explant. In addition, counting ROIs were placed in the mid-region of the BP along the neural to abneural axis to avoid any confounding measurements due to radial differences between tall and short hair cells. For each sample, hair cells were counted in four separate ROIs for each position along the BP. Lumenal surface areas were determined by measuring the circumference of individual hair cells at the level of the cuticular plate. Nuclear size was determined using the DAPI signal.

### Statistical testing and analyses

All data were assessed for normality prior to application of statistical tests, with a threshold of p < 0.05 used for determining significance. When comparing between proximal and distal regions within the same tissue explant, paired t-tests with unequal variance were used. This statistical approach was chosen given that measurements were made from different regions within the same sample and were therefore not independent from each other. Comparisons made between different developmental stages were assumed independent from one another and thus here, independent t-tests and 2-way ANOVAs were used.

### *In situ* hybridisation

Inner ear tissue was dissected and fixed in 4% PFA overnight at 4 °C. Tissue was subsequently washed three times for 30 min in 0.1 M PBS, dehydrated in ascending methanol series (25–100%) and stored at −20 °C until use. Immediately before the *in-situ* protocol, tissue was rehydrated in a descending methanol series (100–25%). Antisense digoxigenin-labelled RNA probes for *Bmp7* were kindly provided by Doris Wu (NIDCD, NIH). *Chd-l1* was synthesised as described previously ^18^. *In situ* hybridisation was performed at 68°C following the protocol as described previously ^66^.

### RNAscope

Gene-specific probes and the RNAscope^®^ Fluorescent Multiplex Reagent Kit (320850) were ordered form Advanced Cell Diagnostics. BP were collected from E8-E10 chick embryos, fixed overnight in 4% paraformaldehyde, and subsequently cryopreserved through a sucrose gradient (5%, 10%, 15%, 20%, and 30%). Samples were embedded in cryomolds using Tissue-Tek O.C.T compound and sectioned on a cryostat at 10-12 μm thickness. RNAscope hybridisation protocol was carried out based on the manufacturer’s (ACD) suggestions. All fluorescent images were obtained on a Zeiss LSM900 confocal microscope.

### RNA-seq analysis

For bulk RNA-seq analysis, all genes with a Log_2_ P-value > 1 were considered significantly expressed in the distal BP region and all genes with a Log_2_ < 1 significantly expressed in the proximal BP region. Statistical significance levels were calculated by one-way ANOVA. For a gene to be considered ‘differential’, at least one region of the BP (proximal, middle or distal) was required to be ≥ 0.5 RPKM. A fold change of ≥ 2 was imposed for the comparison between distal and proximal regions. A final requirement was that middle region samples had to exhibit RPKM values mid-way between proximal and distal regions to selectively capture transcripts with a gradient between the two ends. Bulk Affymetrix data were analysed for differentially expressed mRNAs encoding metabolic effector proteins that regulate cellular NADPH levels. Microarray signals were normalised using the RMA algorithm. The mRNAs expressed at significantly different levels in distal versus proximal BP were selected based on ANOVA analysis using the Partek Genomics Suite software package (Partek, St. Charles, MO, USA). **\*** = p < 0.05.

For detailed description of analysis and protocols please refer to Mann at al., 2014.

## Acknowledgements

The authors wish to thank Dan Jagger, Matthew Kelley, Jeremy Green, Michael Duchen, Gyorgy Szabadkai and Thomas Coate for providing important comments and critical discussion on the manuscript. This project was supported by funds from the King’s Prize Fellowship (King’s College London) (ZFM) and the Biotechnology and Biological Sciences Research Council (BBSRC) grant BB/V006371/1 (ZFM).

## Author contributions

ZFM, TSB conceptualised the study and experimental design.

ZFM, TSB, JO, CS MA developed and executed the methodology.

ZFM, JO, CS, VY analysed and interpreted the data.

ZFM, TSB, JO wrote and edited the manuscript, which was reviewed by all contributing authors.

ZFM is the guarantor of this study, with responsibility for integrity and accuracy of the data.

---

**Figure 2 Supplement 1.**
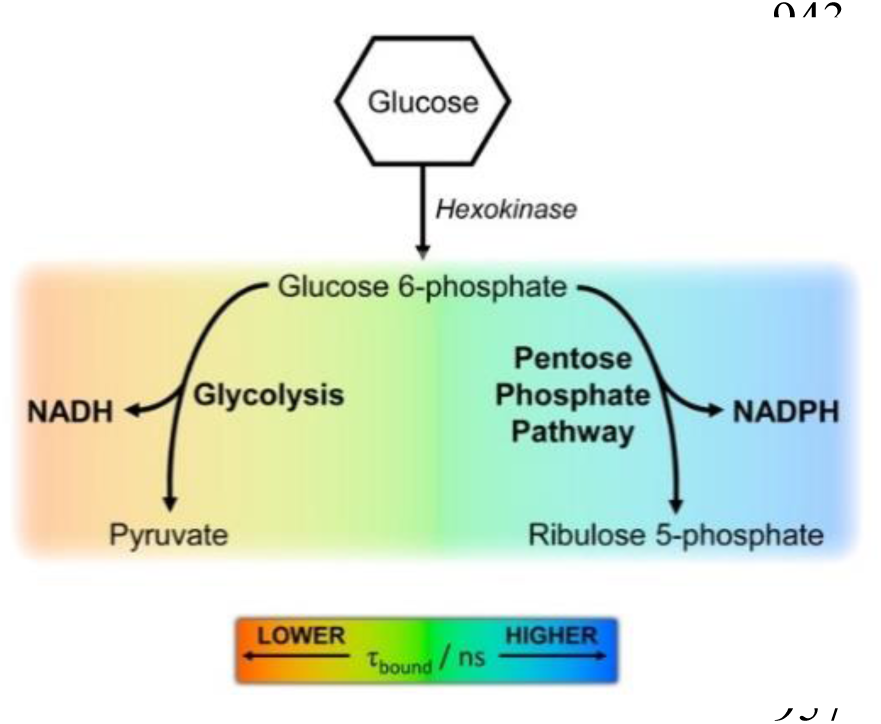
Interpretation of differences in the τ_bound_ signal reported by NAD(P)H FLIM. The gradient in τ_bound_ duration indicates differences in pathways of glucose catabolism. Short lifetimes (orange) indicate NADH production and therefore glucose flux through the main glycolytic pathway, and longer lifetimes (blue) indicate NADPH production and glucose catabolism in the pentose phosphate pathway (PPP). Differences in the τ_bound_ lifetime duration thereby confer differences in cellular metabolic state.

**Figure 3 Supplement 1.**
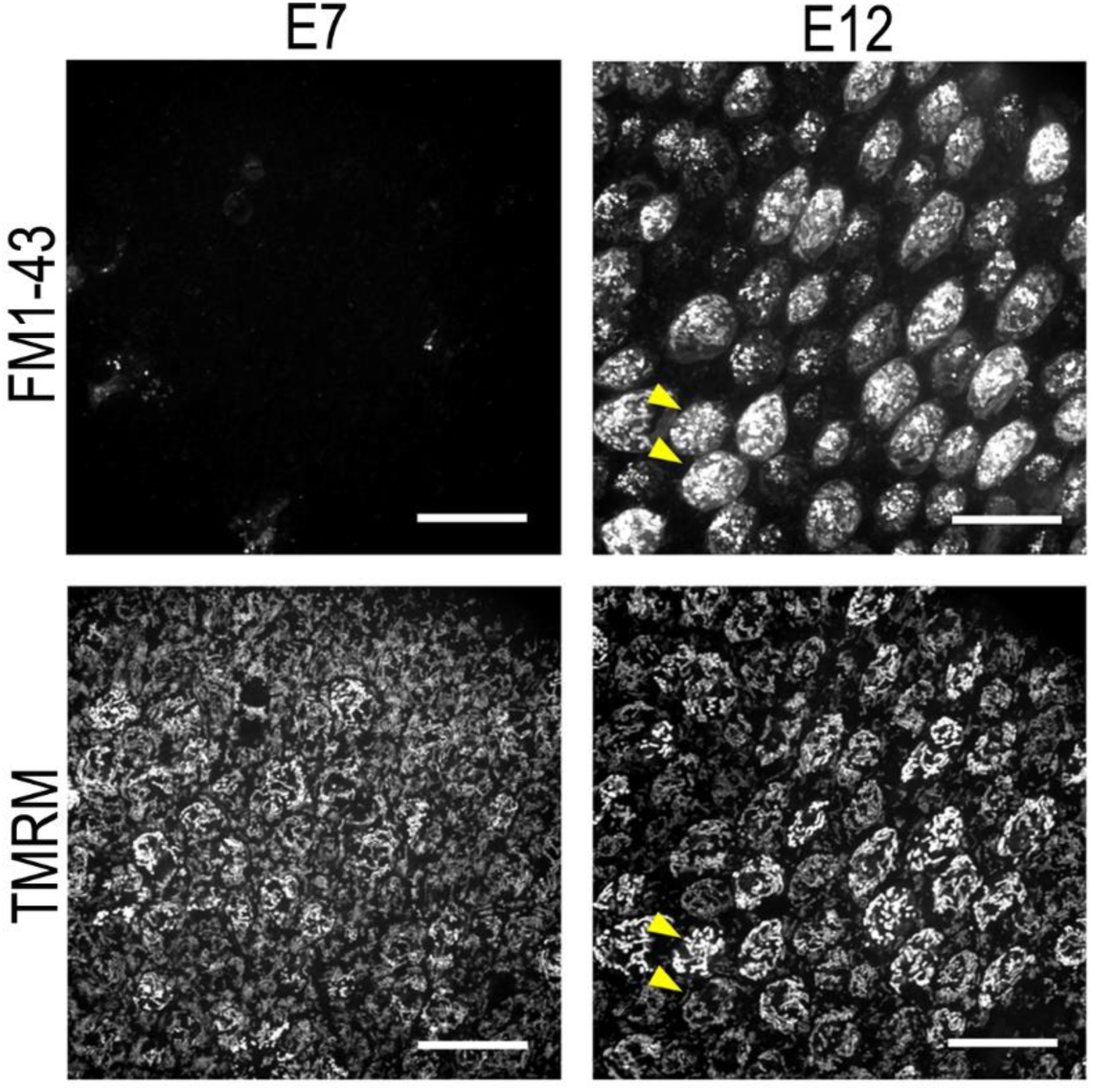
TMRM fluorescence intensity is not dependent on uptake via the HC transduction channel. Live BP explants at E7 and E12 dual loaded with the permeant MET channel blocker FM1-43 (top) and the mitochondrial membrane potential dye TMRM (bottom). FM1-43 provides a measure of HC transduction channel activity. Yellow arrows show that the differences in TMRM fluorescence do not arise from variation in HC transduction channel activity. Scale bars are 20μm.

**Figure 3. Supplement 2.**
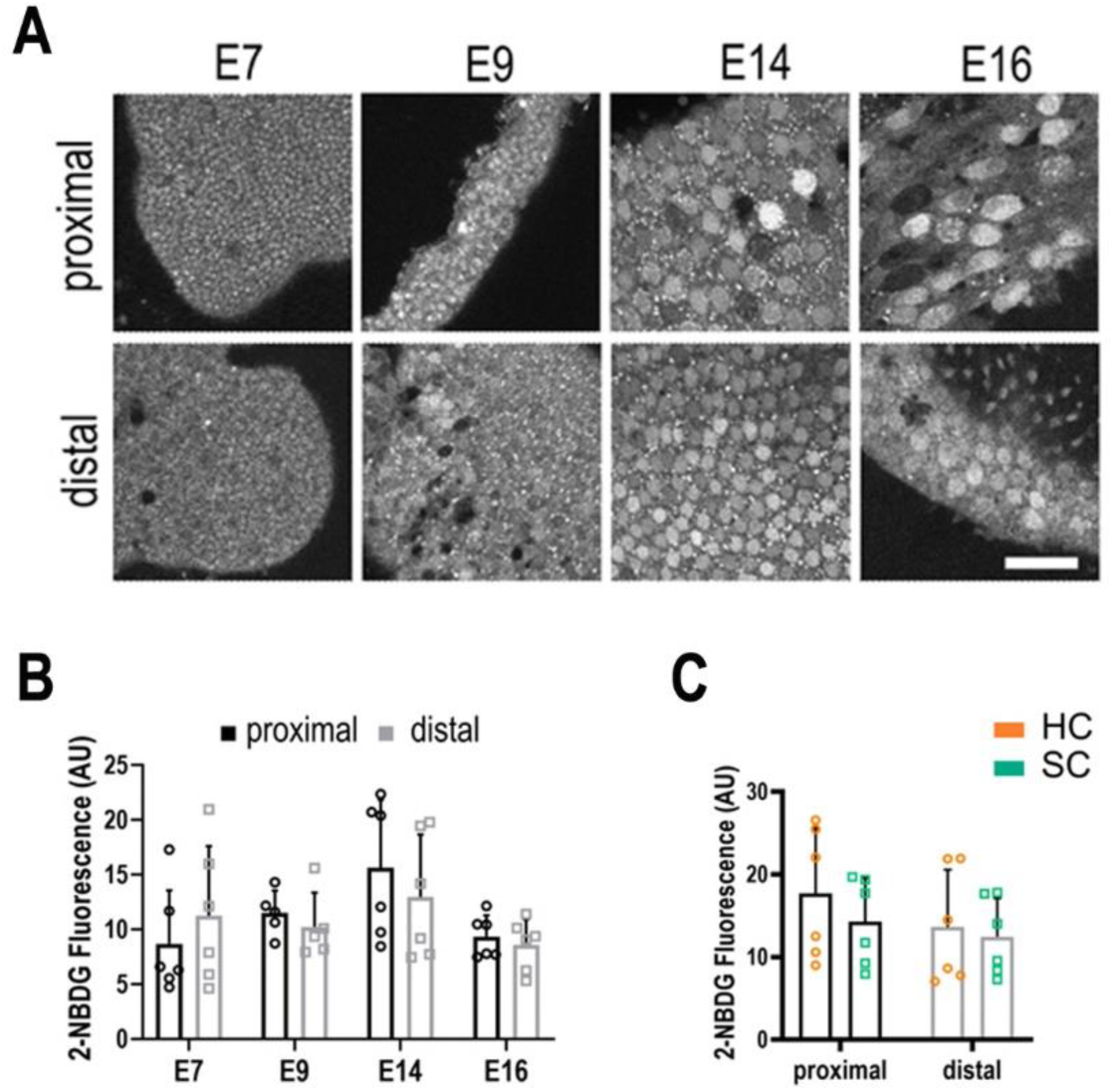
Live imaging of glucose uptake in HCs and SCs at different positions along the tonotopic axis. **(A)** 2-NBDG labelling in live BP explants shows no difference in glucose uptake along the proximal-to-distal axis or between cell types during development. 2-NBDG is a fluorescent glucose analogue which enters cells via GLUT transporters in the plasma membrane and is then phosphorylated by Hexokinase (HK) the first rate-limiting step in glycolysis (Figure 1). 2-NBDG fluorescence therefore reflects the rate of cellular glucose uptake, which can be correlated with its consumption. For instance, tumour cells with elevated glucose metabolism via the Warburg effect exhibit elevated 2-NBDG uptake ^28^. **(B)** Quantification of glucose uptake during development measured from the same explants as indicated in Figure 3. **(C)** 2-NBDG fluorescence in hair cells (orange bars) and supporting cells (green bars) at E14. Data are mean ± SEM. p > 0.05 for proximal and distal regions 2-way ANOVA. E7 n=6, E9 n=5, E14 n=6, E16 n=6 biological replicates. HCs vs SCs n = 6 biological replicates p > 0.05 2-way ANOVA. Scale bars = 40 μm.

**Figure 3 supplement 3.**
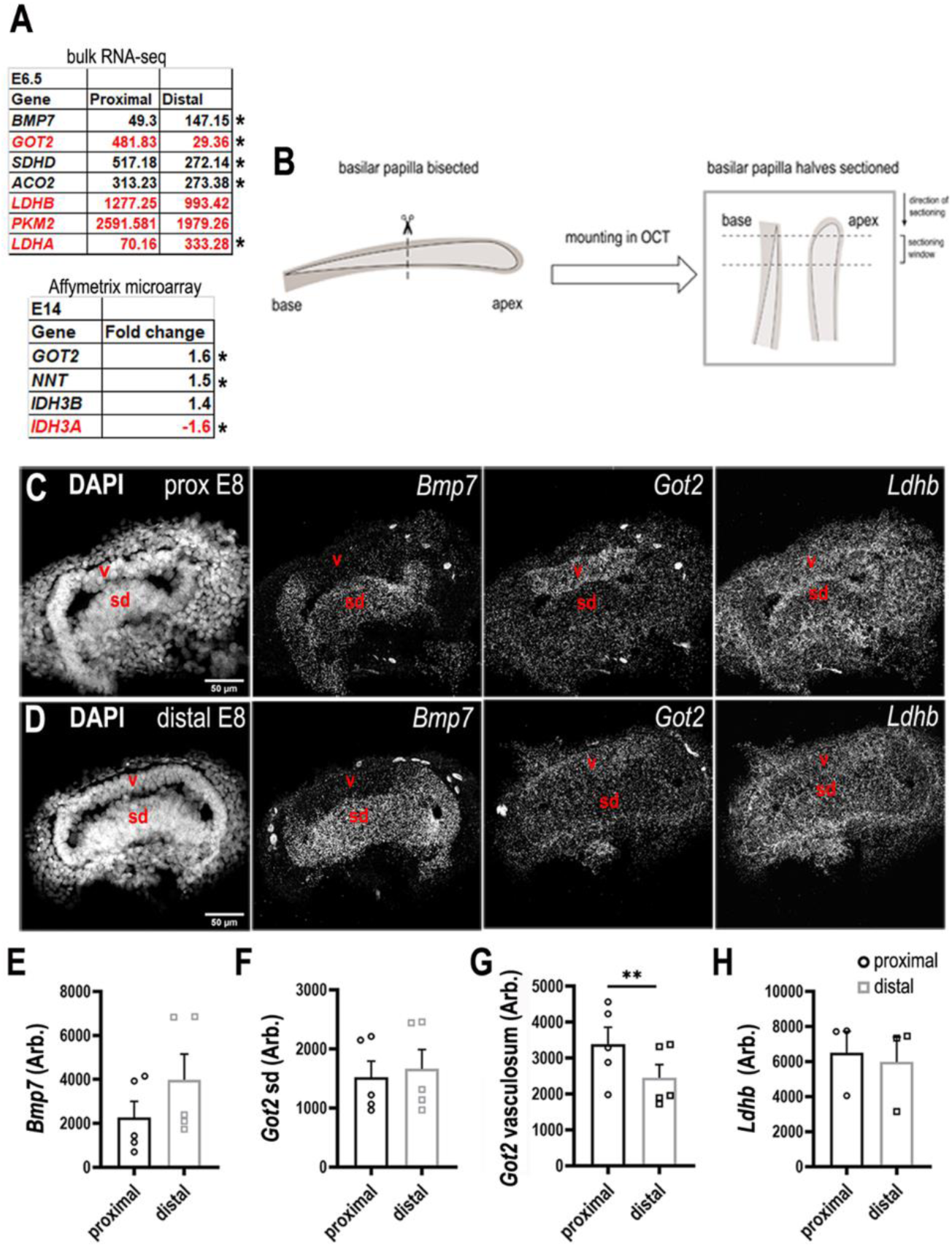
Differential expression of metabolic enzymes along the tonotopic axis of the chick BP. **(A Top)** Metabolic genes with differential expression between proximal and distal BP regions. Data show normalised RPKM values in the proximal and distal BP regions at E6. **(A bottom)** Affymetrix microarray data showing the fold change in expression of each transcript at the proximal compared to the distal region of the BP at E14. **(B)** Sectioning protocol used to collect samples from proximal and distal regions at E8. Proximal and distal tissue used for RNA scope was collected and processed simultaneously from within the same BP. **(C-D)** RNA scope analysis of genes encoding metabolic regulatory proteins in the developing BP identified in A. Images show gene expression in comparative cross sections from proximal and distal BP regions at E8. *Bmp7* expression was used as a control for tonotopic identity as we know it is expressed in a distal-to-proximal gradient. **(E-H)** Quantification of RNA scope fluorescence in the BP at proximal and distal regions. The sensory domain (sd) and tegmentum vasculosum (v) were identified morphologically using the DAPI channel. *Bmp7* – Bone morphogenetic protein 7, *Got2* – Glutamic-Oxaloacetic Transaminase 2, *Sdhd* – Succinate dehydrogenase complex subunit D, *Aco2* – Aconitase 2 *Ldhb* – Lactate dehydrogenase beta, *Pkm2* – Pyruvate kinase M2, *Ldha* – Lactate dehydrogenase alpha, *Nnt* - Nicotinamide Nucleotide Transhydrogenase, *Idh3b* – Isocitrate dehydrogenase beta, *Idh3a* – Isocitrate dehydrogenase alpha. Transcripts annotated red indicates those involved in regulating cellular NADPH/NADH. Data are mean ± SEM, ** = p < 0.001. 2-way ANOVA.

**Figure 3 Supplement 4.**
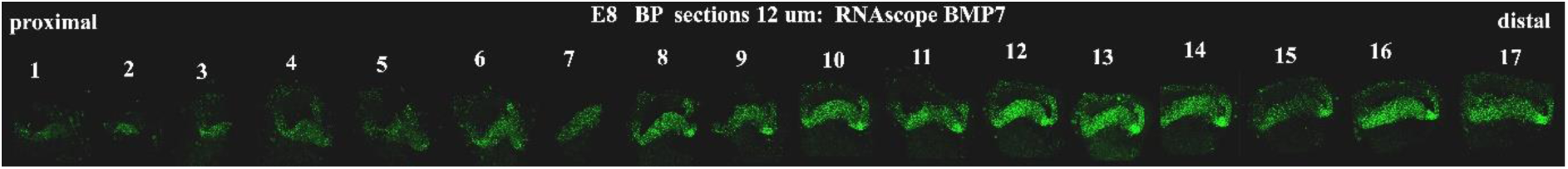
Tonotopic expression of *Bmp7* along the developing BP. RNA scope anlysis of *Bmp7* in cross sections show tonotopic expression along the proximal-to-distal axis at E8.

**Figure 3 Supplement 5.**
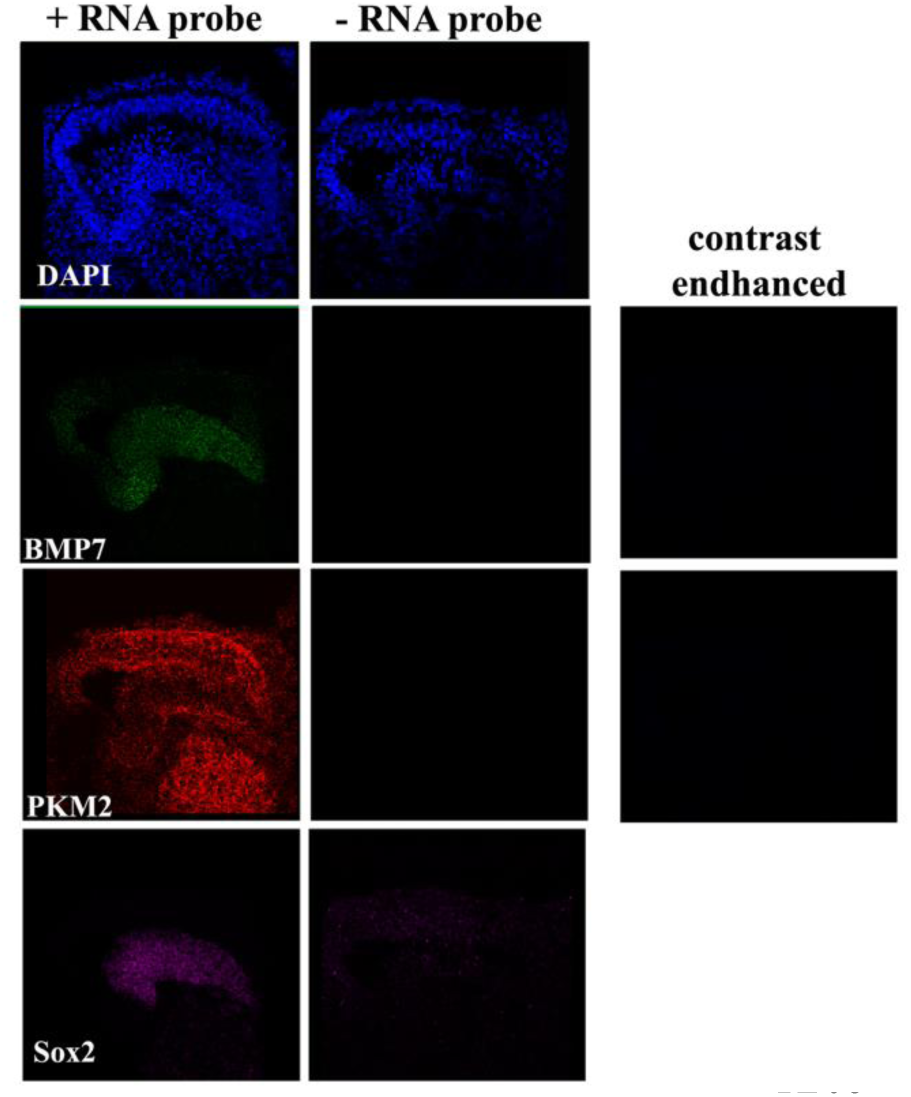
Negative controls for RNA scope analysis. Images show the RNA scope signal in BP cross-sections in the presence and absence of RNA scope probes for *Bmp7*, *Pkm2* and *Sox2*.

**Figure 4 Supplement 1.**
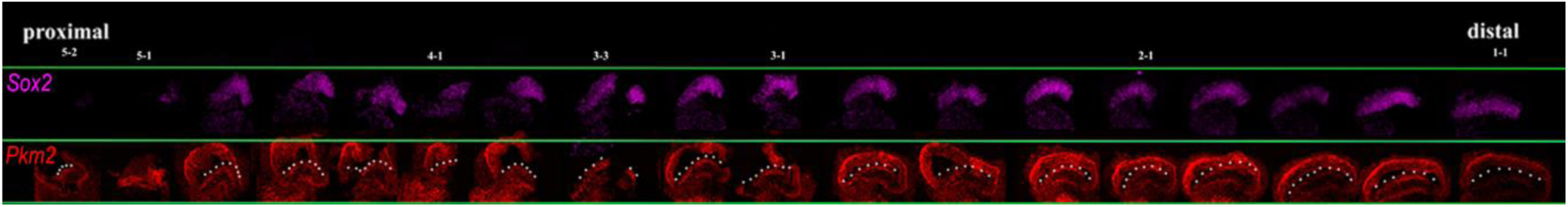
Proximal-to-distal expression of *Sox2* and *Pkm2* at E8. Images show cross-sections of the chick BP at E8 indicating expression of *Sox2* and *Pkm2* along the proximal-to-distal axis. White dashed line indicates the prosensory region.

**Figure 4 Supplement 2.**
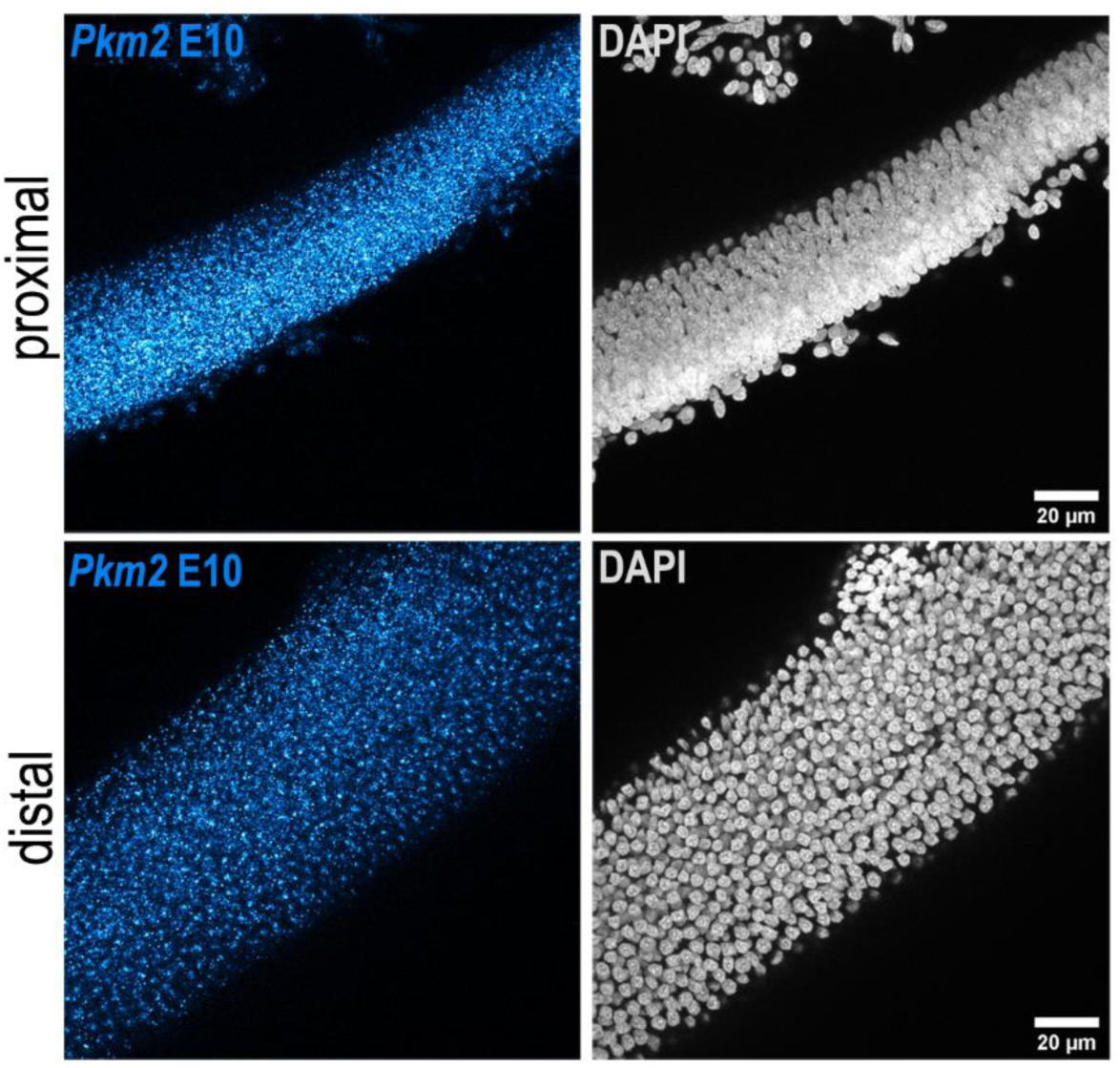
Tonotopic expression of *Pkm2* in the developing BP. RNA scope analysis of *Pkm2* expression in BP whole-mounts from the proximal and distal regions at E10. Data are representative of 4 independent biological replicates.

**Figure 4 Supplement 3.**
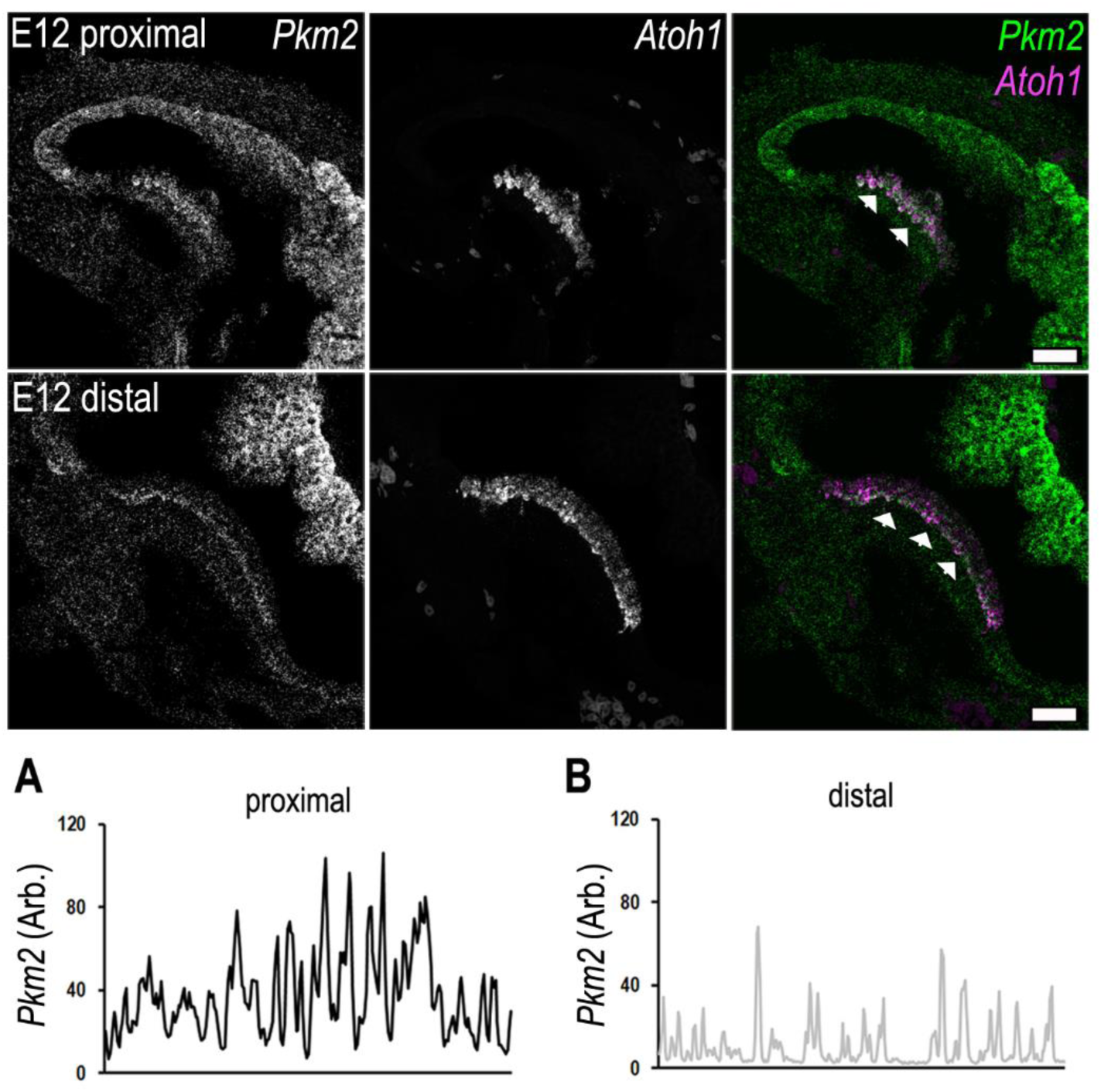
Tonotopic expression of *Pkm2* and *Atoh1* in the developing BP. RNA scope analysis of *Pkm2* and the hair cell-specific transcription factor *Atoh1* in BP whole-mounts of the proximal and distal regions at E12. Data are representative of 3 independent biological replicates. **(A-B)** Line scan analysis along the prosensory region in the proximal and distal regions highlighting differences in *Pkm2* fluorescence intensity. Scale bar is 20 μm. Data are representative of 3 biological replicates.

**Figure 5 Supplement 1.**
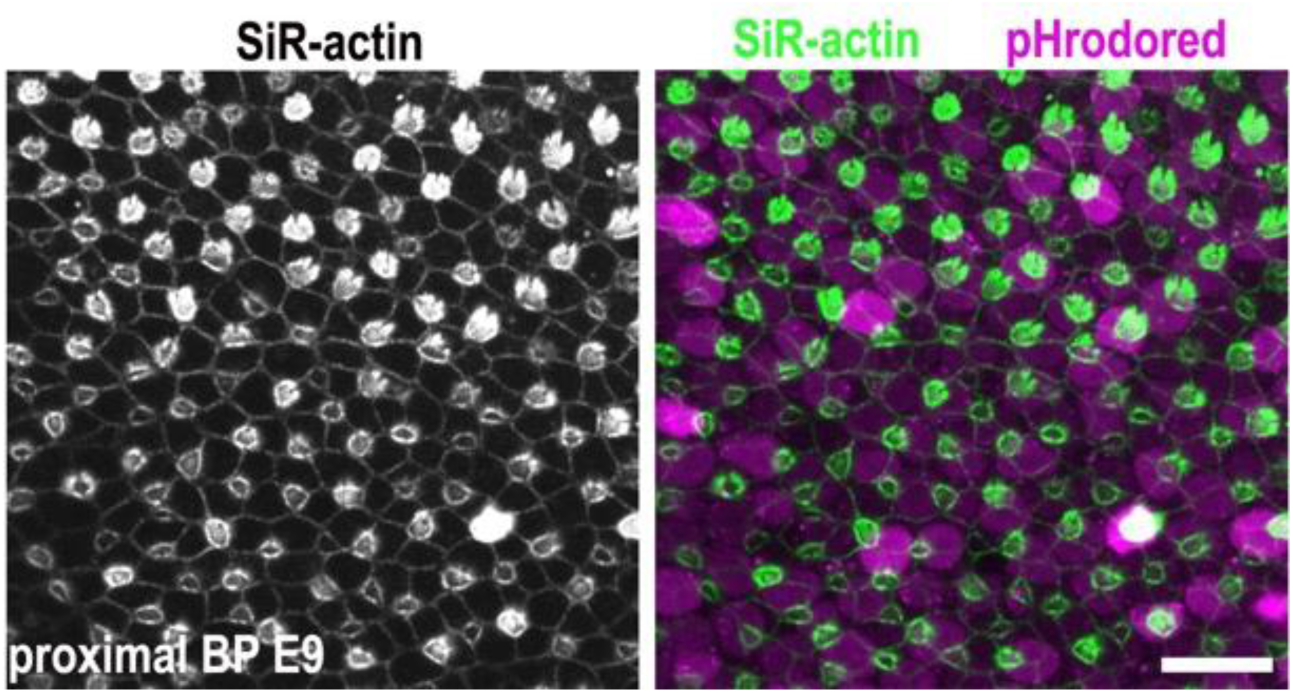
Generating HC and SC analysis masks. Images show z-projections of BP explants dual loaded with pHrodo Red and SiR-actin. The stereocilial bundles at the epithelial surface were used to differentiate between HCs and SCs when generating analysis masks.

**Figure 5 Supplement 2.**
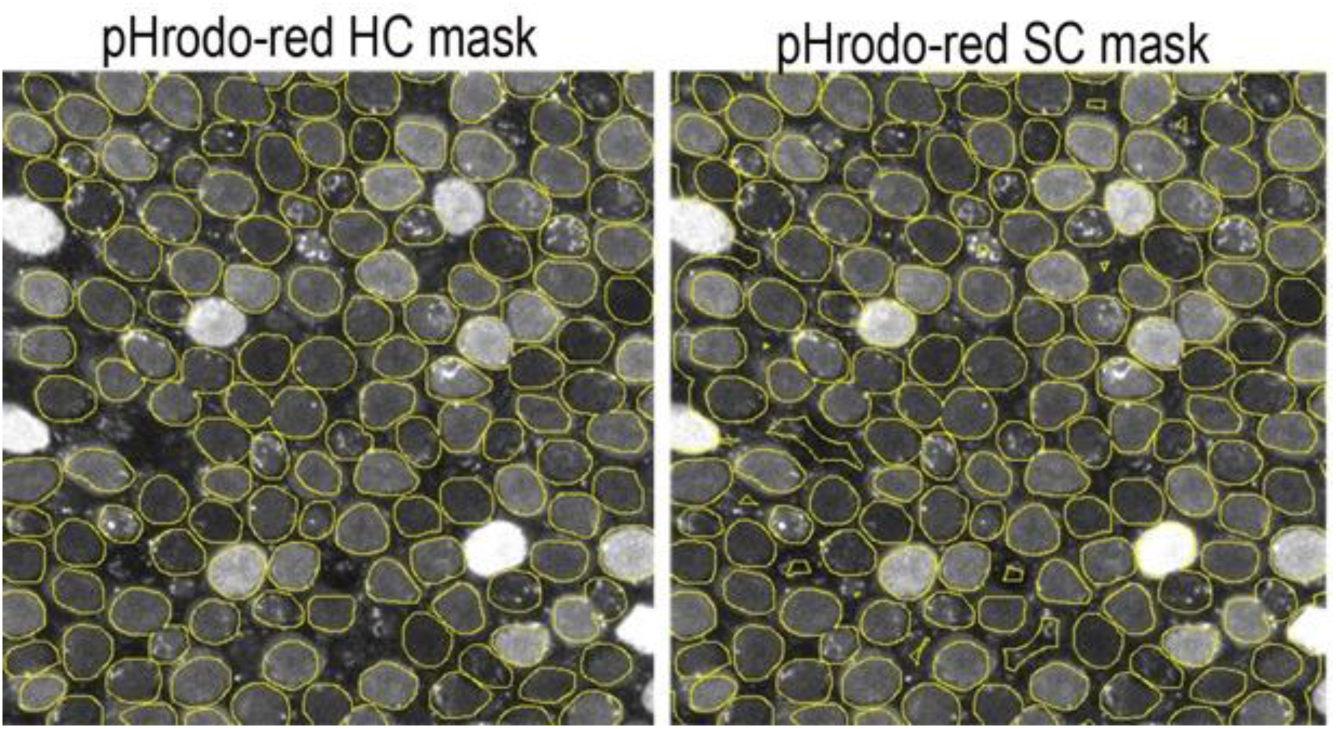
HC and SC masks used for quantification of pHrodo Red fluorescence. HC fluorescence was measured from within the ROIs (yellow circles). SC fluorescence was measured from inverse masks generated using the HC ROIs. SC signal was assumed to originate from the areas outside of the defined HC ROIs. The presence of a stereociliary bundle as reported by the SiR-actin was used to differentiate between HCs and SCs.

**Figure 5 Supplement 3.**
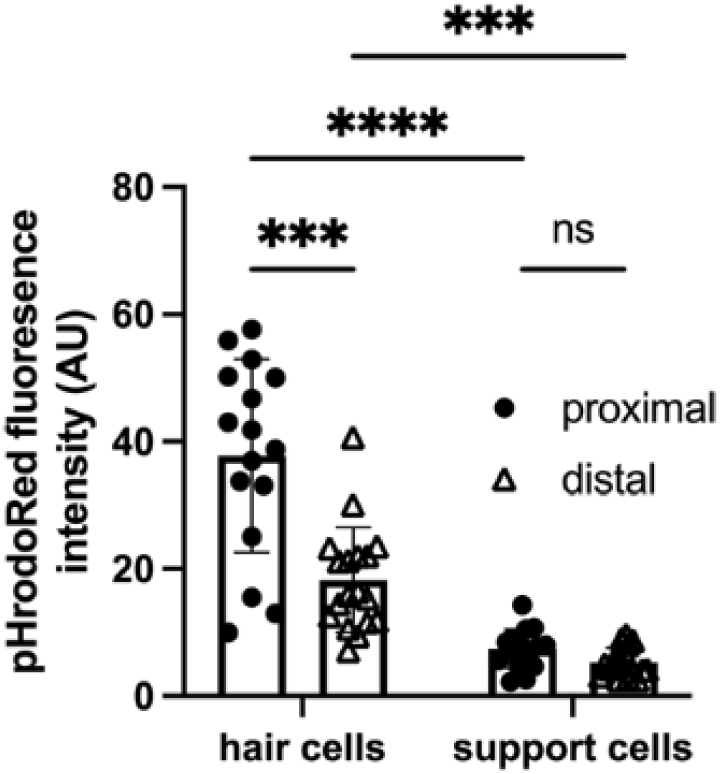
pHrodo Red fluorescence along the tonotopic axis of the chick BP at E14. Quantification of mean pHrodo Red fluorescence in HCs and SCs from proximal and distal BP regions at E14 shows a reverse gradient to that reported by the indicator at early developmental stages. Data are mean ± SEM from 16 independent biological replicates. * p = < 0.05, *** p = < 0.001, **** p = < 0.0001 2-way ANOVA.

**Figure 6 Supplement 1.**
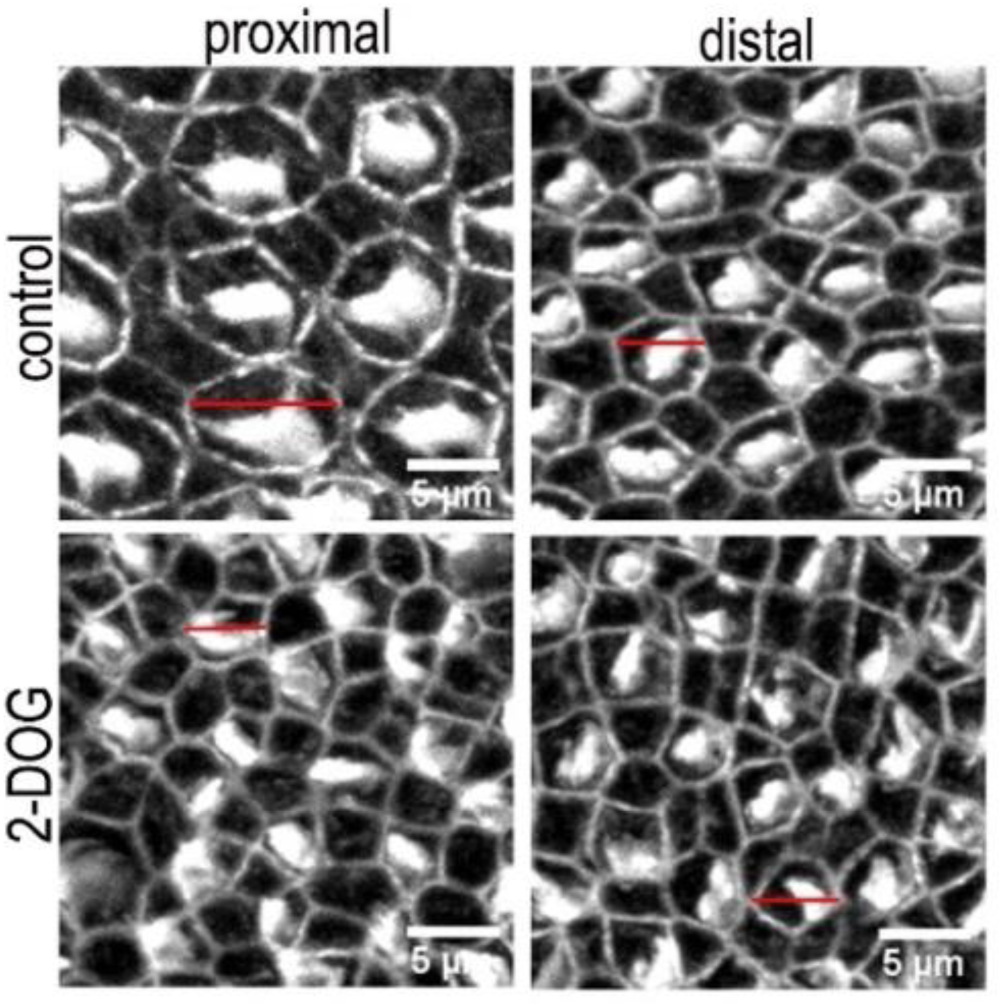
Morphological differences in the HC bundle and cuticular plate region in BP explants treated with control or 2-DOG containing medium. Images are maximum z-projections of Phalloidin stained BP explants from the proximal and distal BP regions. Explants were maintained from E6.5 for 7 days *in vitro* (equivalent to E13.5) in either control medium or that supplemented with 2 mM 2-DOG + 5 mM Sodium Pyruvate (NaP). Phalloidin staining depicts differences in hair cell lumenal surface area and gross bundle morphology between proximal and distal regions. Red lines indicate the size difference in HC lumenal surface area between control and 2-DOG-treared cultures.

**Figure 6 supplement 2.**
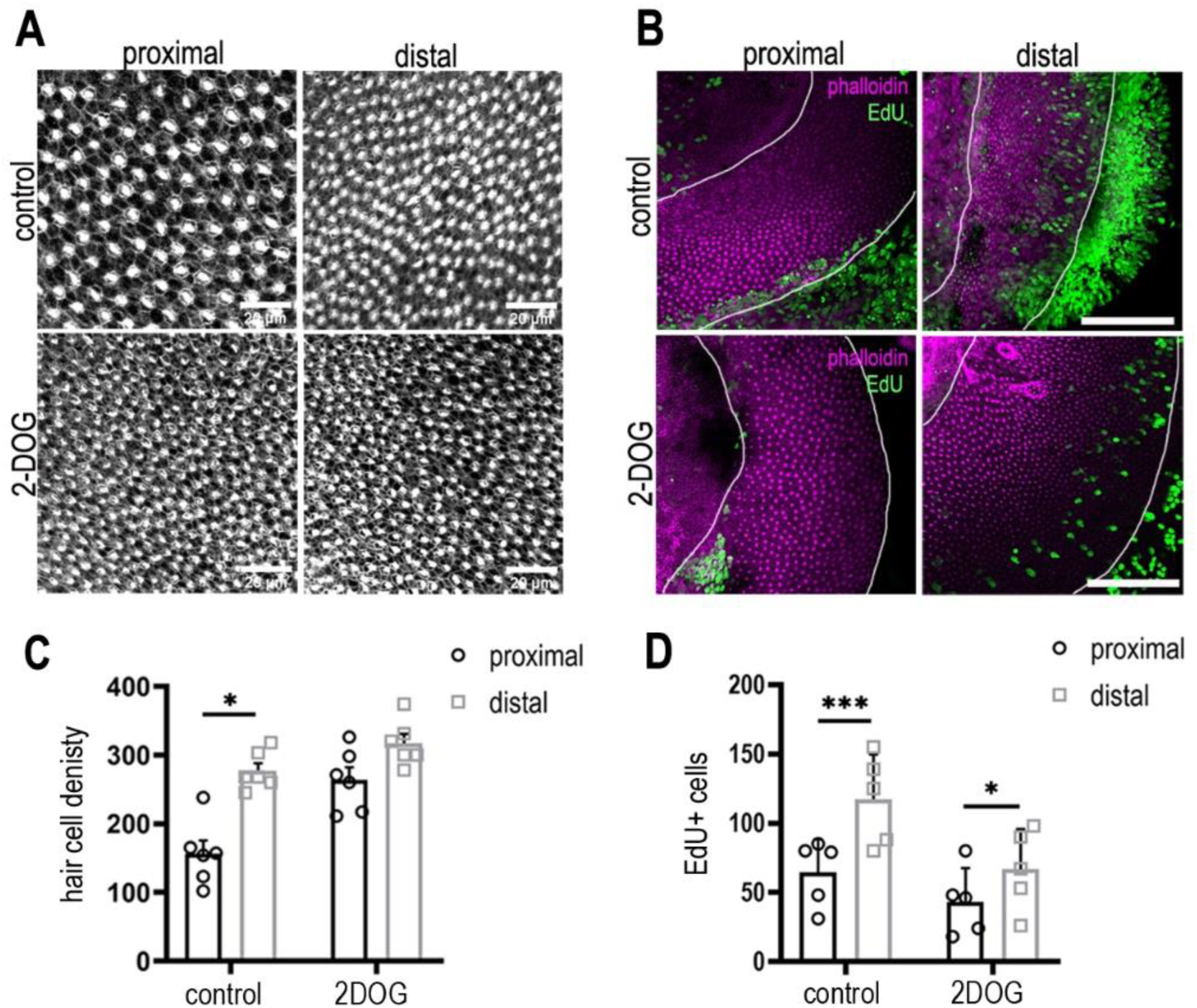
2-DOG increases HC cell density in the proximal BP region independently of proliferation. (A) Phalloidin staining indicating HC density in the proximal and distal BP regions in control (top) and 2-DOG treated (bottom) explants. **(B)** Maximum z-projections of proximal and distal regions from Phalloidin (magenta) and the EdU (green) stained BPs. Explant cultures were established at E8 and maintained for 48 hours *in vitro* in control medium +EdU or medium containing EdU + 2 mM 2-DOG and 5 mM sodium pyruvate. Proliferation was consistently reduced in 2-DOG treated explants. **(C)** Quantification of HC density in proximal and distal BP regions counted from 100 mm^2^ ROIs in each region. **(D)** EdU counts from 100 mm^2^ ROIs in proximal and distal BP regions in control and 2DOG-treated cultures. Data mean ± sem. * = p < 0.05, *** p < 0.001 2-way ANOVA. Control: n = 5 biological replicates, 2-DOG: n = 5 biological replicates. EdU: control n= 5 independent biological replicates; 2-DOG EdU n = 5, 2-DOG HC density n = 6 independent biological replicates. White lines outline the regions of BP sensory epithelium.

**Figure 6 Supplement 3.**
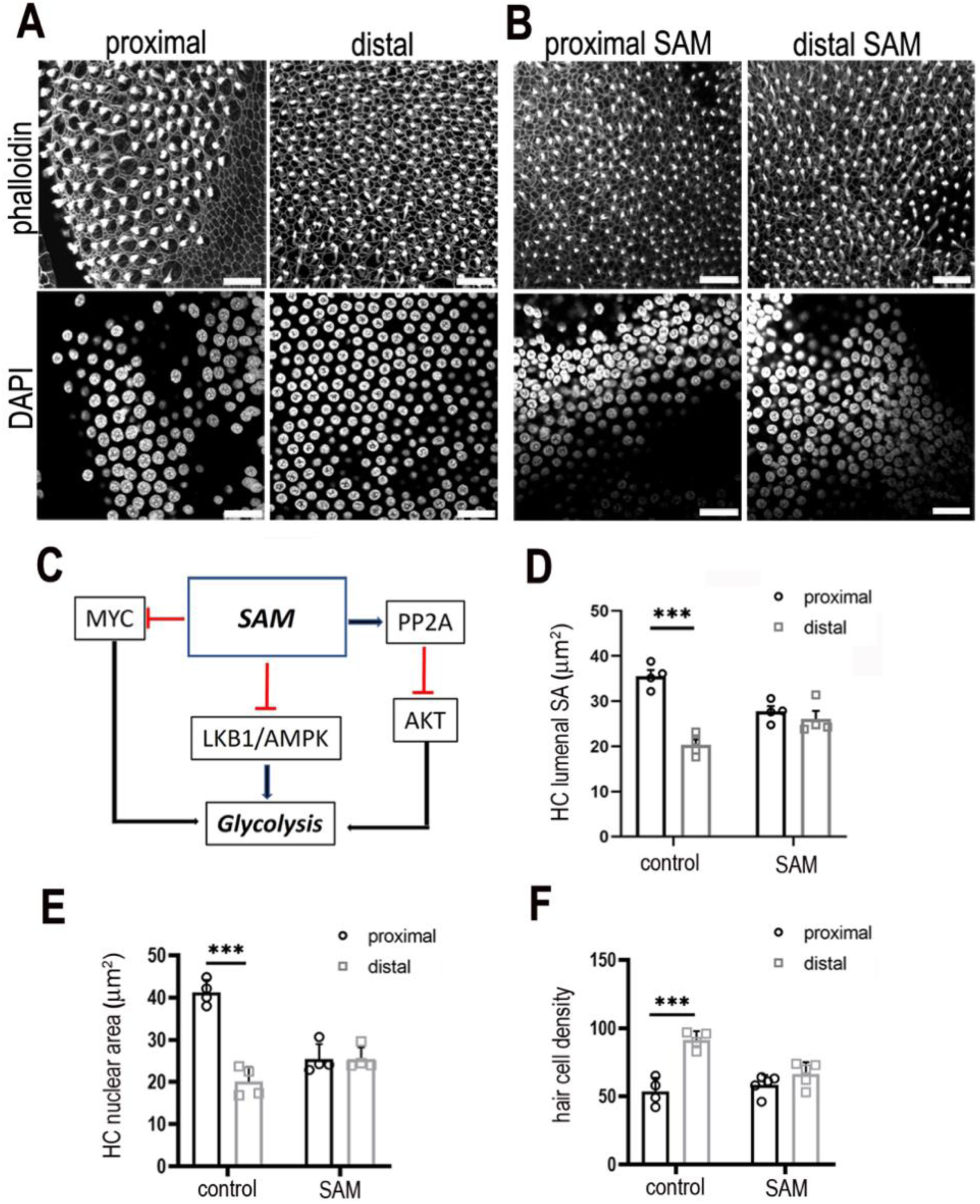
Modulating S-Adenosyl methionine during development abolishes the gradient in hair cell morphology along the tonotopic axis of the BP. **(A-B)** Images are maximal z-projections of proximal and distal regions from BP explants maintained for 7 days *in vitro* in control medium or medium containing 50 μM SAM. Phalloidin staining (A, B top panels) shows the BP surface and hair cell lumenal surface areas and DAPI staining (A, B bottom panels) shows the hair cell size and density along the proximal-to-distal axis. **(C)** Schematic showing how increasing SAM levels indirectly modulates cytosolic glycolytic flux. **(D-F)** Treatment of explants with SAM from E6.5 for 7 days *in vitro* abolished the normal gradient in hair lumenal surface area, nuclear area and cell density along the tonotopic axis. This effect was most pronounced in the proximal compared to distal region. All metrics were quantified from 2500 μm^2^ ROIs in the proximal (black bars) and distal (grey bars) BP regions of control and SAM-treated explants SA = surface area. Data are mean ± SEM. **\*\*\*** p = < 0.001 2-way ANOVA. Controls: n = 4 and SAM: n = 4 biological replicates. Scale bars are 20 μm. MYC - MYC Proto-Oncogene (basic helix-loop-helix transcription factor), SAM - S-Adenosyl methionine, PP2A – protein phosphatase 2, AKT - *Akt* serine/threonine kinase family, LKB1/AMPK – serine threonine kinase pathway in volved in metabolic regulation.

**Figure 7 Supplement 1.**
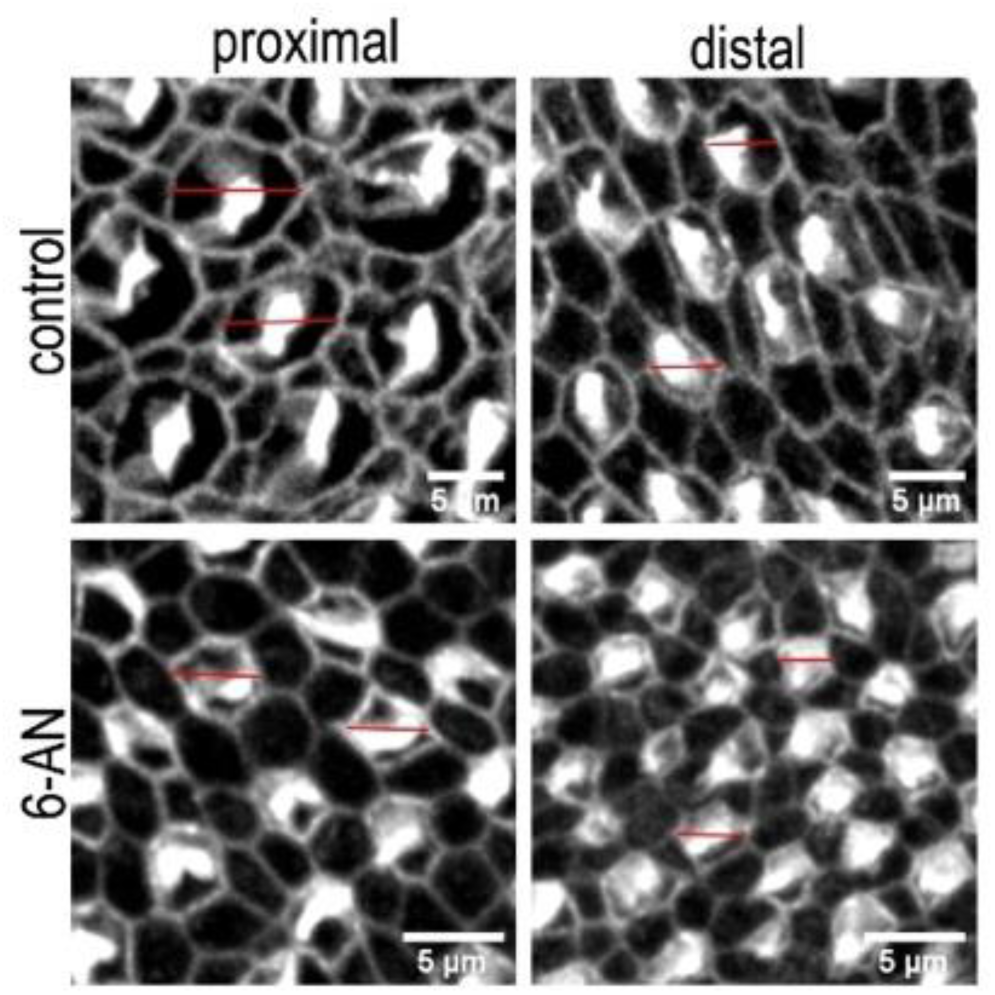
Differences in the HC lumenal surface area and gross bundle morphology in proximal and distal BP regions from explants treated with control or 6-AN containing medium. Images are maximum z-projections of Phalloidin stained BP explants from the proximal and distal BP regions. Explants were maintained from E6.5 for 7 days *in vitro* (equivalent to E13.5) in either control medium or that supplemented with the PPP inhibitor 6-AN. Phalloidin staining depicts differences in hair cell lumenal surface area and gross bundle morphology between proximal and distal regions. Red lines indicate the size difference in HC lumenal surface area between control and 6-AN-treated cultures. Scale bars are 5 μm.

**Figure 7 Supplement 2.**
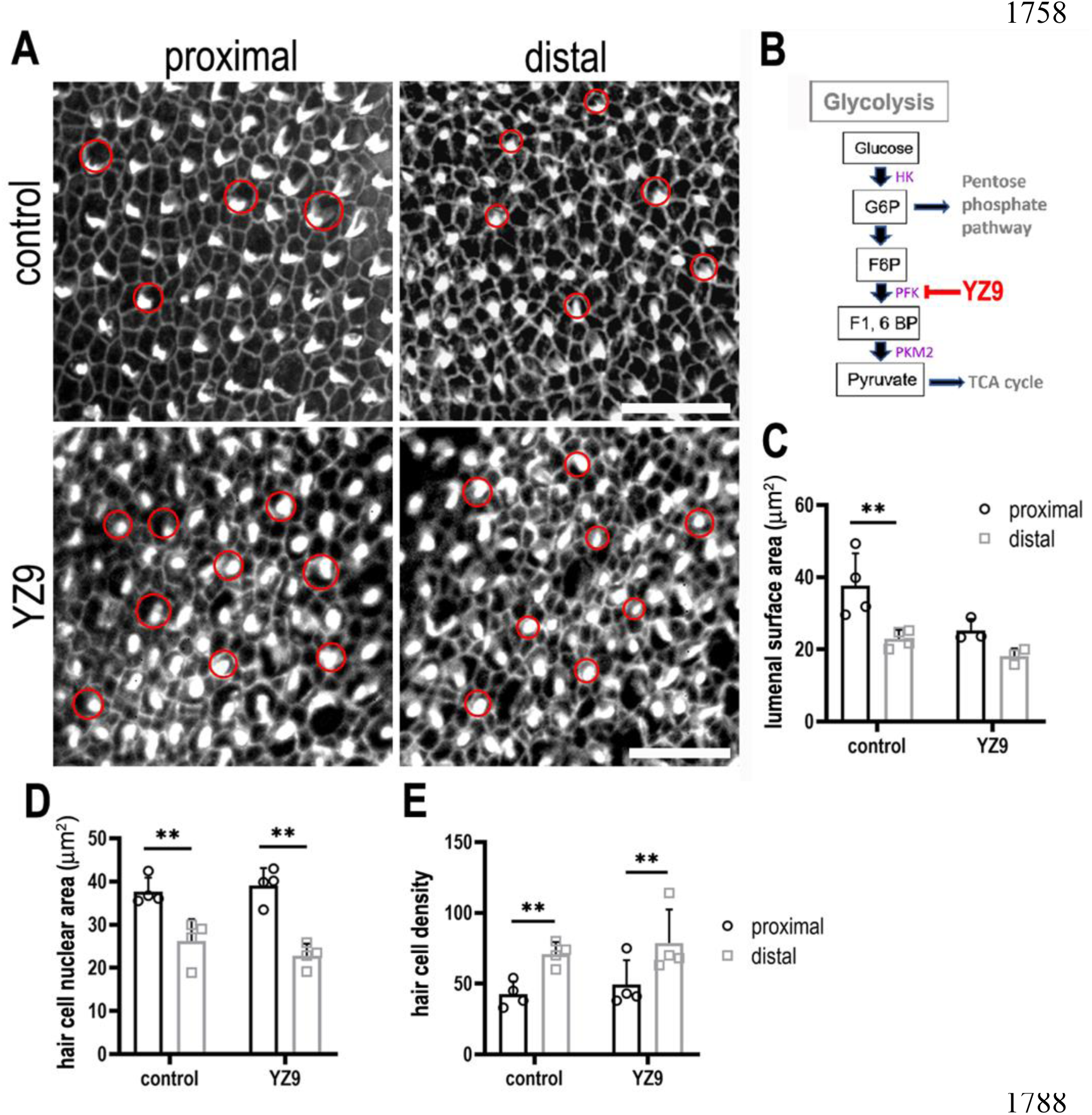
Inhibiting phosphofructokinase alters HC morphology at the luminal surface area during BP development. **(A)** Maximal z-projections showing Phalloidin staining in proximal and distal BP regions of control and YZ9-treated explants. **(B)** Blocking glycolysis at the level of PFK between E6.5 and E13.5 had no effect on HC morphology along the tonotopic axis. Hair cell circumference was quantified in 2500 μm^2^ areas in proximal (black bars) and distal (grey bars) regions, showing no difference in size or density after treatment with 1 μM YZ9. **(C)** Schematic illustrating the inhibitory action of YZ9 in the glycolytic pathway. Data are mean ± SEM. * p < 0.05, ** p = < 0.01 2-way ANOVA. Controls: n = 4, YZ9: n = 4. Scale bars are 20 μm. Red circles indicate size hair cell lumenal surface area between control and YZ9-treated explants (n = 3). HK-Hexokinase, G6P – glucose 6-phosphate, F6P – fructose 6-phosphate, F1, PFK-phosphofuctokinase, 6BP – fructose 1,6-bisphosphate, PKM2 – pyruvate kinase M2.

**Figure 9 Supplement 1.**
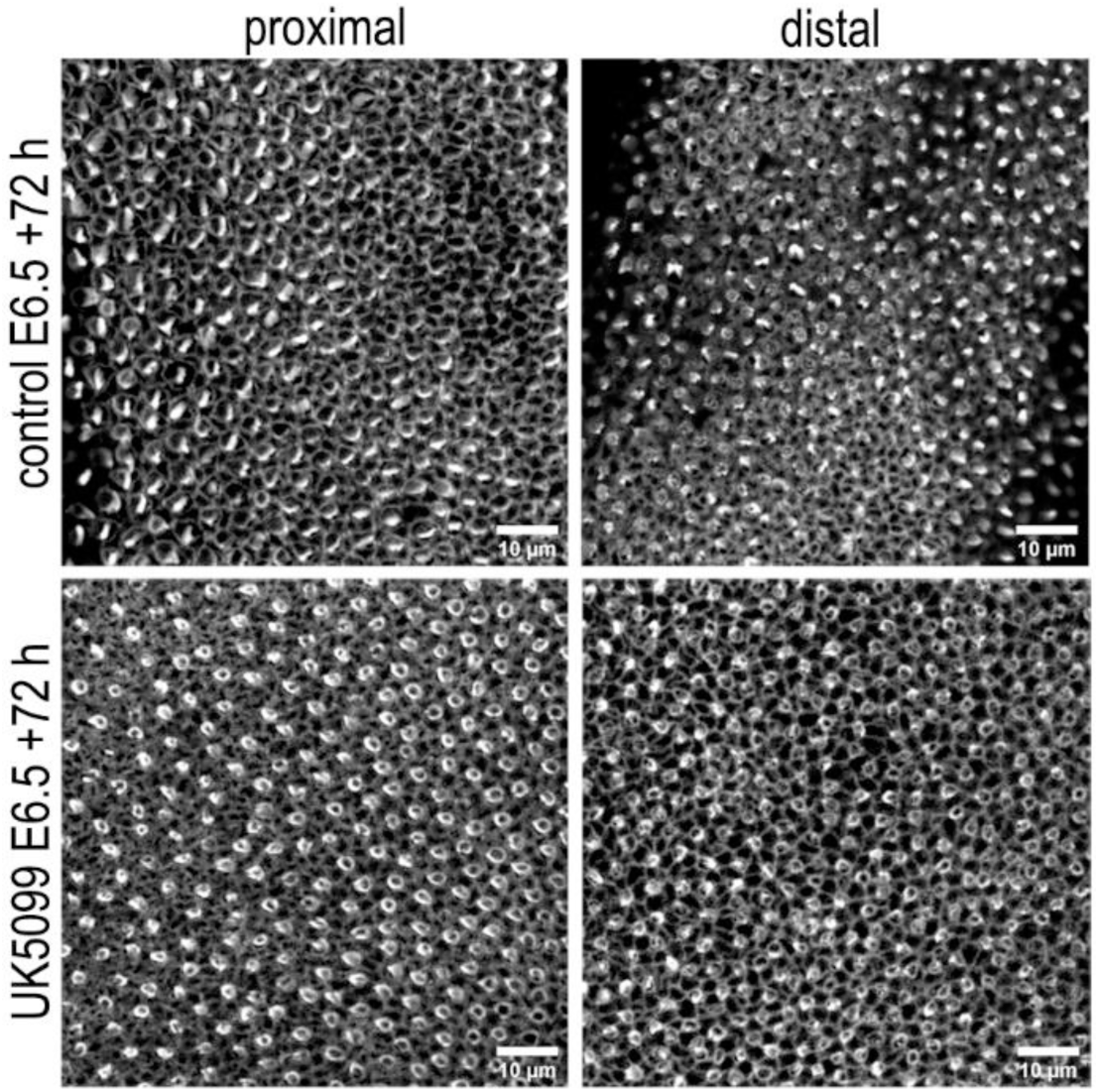
Blocking mitochondrial OXPHOS delays developmental acquisition of normal HC morphologies. Images show HC morphology at the surface of the BP epithelium in explants stained with Phalloidin. Explants were established at E6.5 and maintained for 72 hours (E9.5 equivalent) in culture in control (top) medium or that supplemented with the mitochondrial inhibitor UK5099 (bottom). Images are representative of 3 independent biological replicates.

**Figure 10 Supplement 1.**
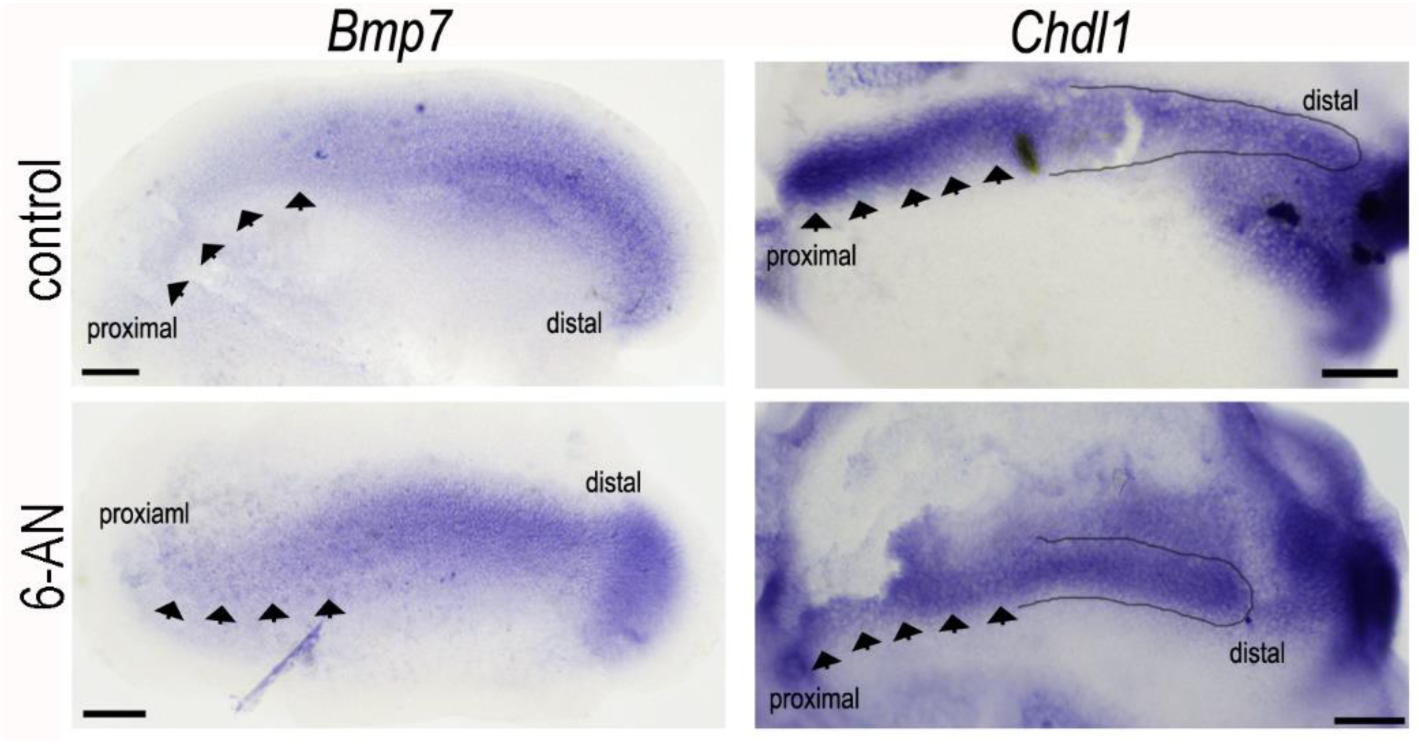
**Blocking glucose flux into the PPP with 6-AN alters the normal expression of *Bmp7* and *Chdl1* in the developing BP**. Treatment of explant cultures with 6-AN from E6.5 for 72 hours *in vitro* caused marked changes in the expression levels of *Bmp7* and *Chdl1*. 6-AN induced a slight expansion of the *Bmp7* gradient into the proximal region (indicated by black arrows) and a reduction in *ChdlI* at the proximal but not distal end (indicated by black arrows and outlined area). Images are representative of 4 independent biological replicates for each probe.

**Figure 10 supplement 2.**
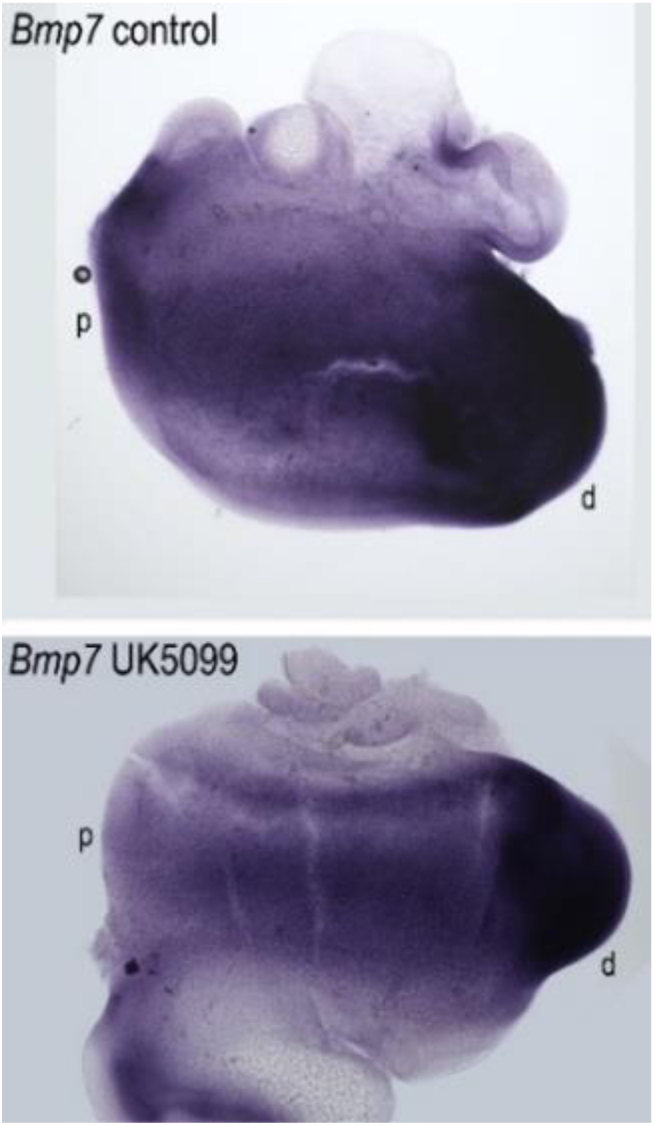
**Blocking mitochondrial metabolism does not alter the gradient of *Bmp7* expression along the developing BP**. Treatment of explant cultures with UK5099 from E6.5 for 72 hours *in vitro* does not elicit marked changes in the normal expression of *Bmp7* along the BP. Images show *in situ* hybridisation for *Bmp7* in BP whole-mounts treated with UK5099 from E6.5 for 72 hours *in vitro*. Images are representative of 5 independent biological replicates.

**Figure 11 Supplement 1.**
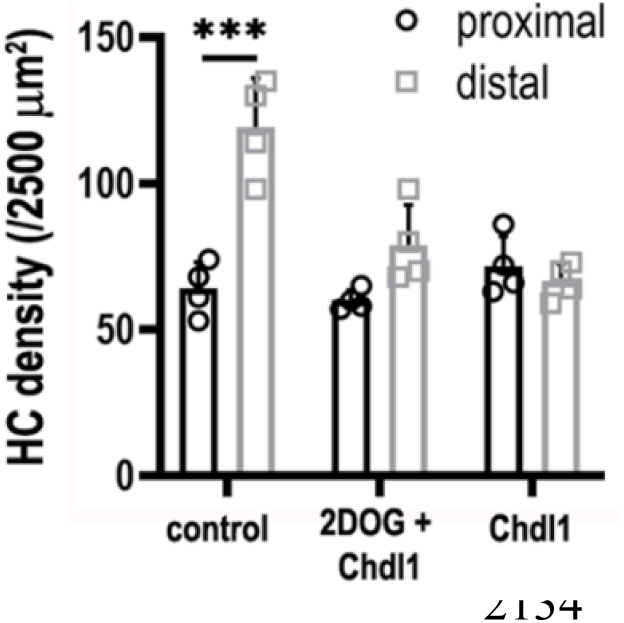
Treatment of explants with 2DOG + Chdl1 partially rescues the pattern of HC density along the developing BP. HC density was quantified in 2500 μm^2^ ROIs in proximal and distal regions. BP explants were established at E6.5 and maintained for 7 days *in vitro* in control conditions, or in medium containing 2-DOG + 0.4 μg/mL Chdl1 or 0.4 μg/mL Chdl1. Data are mean ± SEM from 4 independent biological replicates. *** = p <0.001 2-way ANOVA.

**Supplementary Table 1.** 2-way repeated measures ANOVA results for hair cell nuclear area. p-values in the left-hand column indicate whether the proximal-distal differences in HC nuclear area was altered in inhibitor treated explants compared to controls. The p-values in the *interaction* column indicate whether the proximal-distal differences in nuclear area changed after inhibitor treatment. YZ9 and Chdl1 + 2DOG were the only treatments which did not produce a significant interaction, indicating preservation of the tonotopic gradient.

| Treatment | 2-way RM ANOVA p-values |  | significant interaction? |
| --- | --- | --- | --- |
|  | overall proximal-distal differences | interaction |  |
| 6AN | 0.0005 | 0.0002 | *** |
| 2-DOG | 0.0002 | 0.0001 | *** |
| SAM | 0.0011 | 0.0003 | *** |
| YZ9 | 0.0055 | 0.1962 | ns |
| UK-5099 | 0.0287 | 0.0112 | * |
| BMP7 | 0.0001 | 0.0001 | *** |
| Chdl-1 | 0.0001 | 0.0002 | *** |
| Chdl-1+2-DOG | 0.0007 | 0.7342 | ns |
| Shikonin | 0.0002 | 0.0396 | * |
| Shikonin+6AN | 0.0066 | 0.0136 | * |

**Supplementary Table 2.** 2-way repeated measures ANOVA results for hair cell luminal surface area. p-values in the left-hand column indicate whether the proximal-distal differences in HC luminal SA was altered in inhibitor treated explants compared to controls. The p-values in the *interaction* column indicate whether the proximal-distal differences in luminal SA changed after treatment. YZ9 and Chdl1 + 2DOG were the only treatments which did not produce a significant interaction, indicating preservation of the tonotopic gradient.

| Treatment | 2-way RM ANOVA p-values |  | significant interaction? |
| --- | --- | --- | --- |
|  | overall proximal-distal differences | interaction |  |
| 6AN | 0.0005 | 0.0002 | *** |
| 2-DOG | 0.0002 | 0.0001 | *** |
| SAM | 0.0011 | 0.0003 | *** |
| YZ9 | 0.0055 | 0.1962 | ns |
| UK-5099 | 0.0287 | 0.0112 | * |
| BMP7 | 0.0001 | 0.0001 | *** |
| Chdl-1 | 0.0001 | 0.0257 | * |
| Chdl-1+2-DOG | 0.0015 | 0.3702 | ns |
| Shikonin | 0.0021 | 0.0431 | * |
| Shikonin+6AN | 0.0007 | 0.0056 | ** |

